# COQ4 is required for the oxidative decarboxylation of the C1 carbon of Coenzyme Q in eukaryotic cells

**DOI:** 10.1101/2023.11.13.566839

**Authors:** Ludovic Pelosi, Laura Morbiato, Arthur Burgardt, Fiorella Tonello, Abigail K. Bartlett, Rachel M. Guerra, Katayoun Kazemzadeh Ferizhendi, Maria Andrea Desbats, Bérengère Rascalou, Marco Marchi, Luis Vázquez-Fonseca, Caterina Agosto, Giuseppe Zanotti, Morgane Roger-Margueritat, María Alcázar-Fabra, Laura García-Corzo, Ana Sánchez-Cuesta, Plácido Navas, Gloria Brea-Calvo, Eva Trevisson, Volker F. Wendisch, David J. Pagliarini, Leonardo Salviati, Fabien Pierrel

**Author notes:** Address correspondence to Leonardo Salviati or Fabien Pierrel. the authors contributed equally to this article (Ludovic Pelosi, Laura Morbiato).

## Abstract

Coenzyme Q (CoQ) is a redox lipid that fulfills critical functions in cellular bioenergetics and homeostasis. CoQ is synthesized by a multi-step pathway that involves several COQ proteins. Two steps of the eukaryotic pathway, the decarboxylation and hydroxylation of position C1, have remained uncharacterized. Here, we provide evidence that these two reactions occur in a single oxidative decarboxylation step catalyzed by COQ4. We demonstrate that COQ4 complements an *Escherichia coli* strain deficient for C1 decarboxylation and hydroxylation and that COQ4 displays oxidative decarboxylation activity in the non-CoQ producer *Corynebacterium glutamicum*. Overall, our results substantiate that COQ4 contributes to CoQ biosynthesis, not only via its previously proposed structural role, but also via oxidative decarboxylation of CoQ precursors. These findings fill a major gap in the knowledge of eukaryotic CoQ biosynthesis, and shed new light on the pathophysiology of human primary CoQ deficiency due to *COQ4* mutations.

## INTRODUCTION

Coenzyme Q (CoQ) is a redox-active lipid that plays crucial roles in the cellular homeostasis of eukaryotic cells ^1^. CoQ is composed of a quinone head group and a polyisoprenoid chain of variable length: six isoprene units in the yeast *Saccharomyces cerevisiae* (CoQ_6_), ten in humans (CoQ_10_), and eight in the bacterium *Escherichia coli* (CoQ_8_). Despite the crucial functions of CoQ, its biosynthetic pathway in eukaryotes is still incompletely understood. The CoQ biosynthetic machinery includes the products of at least 15 genes (collectively known as *COQ* genes), most of them with a conserved function ^2,3^. Primary CoQ deficiency is a clinically and genetically heterogeneous group of conditions characterized by impairment of CoQ biosynthesis and caused by defects in any of the COQ genes ^4^.

The precursors of CoQ, 4-hydroxybenzoic acid (4-HB) and polyprenyl-pyrophosphate (PPP) are conjugated to form polyprenyl-4-hydroxybenzoic acid (nP-HB). This reaction is catalyzed by COQ2, a prenyltransferase located in the mitochondrial inner membrane ^5^ (Fig. 1A). The subsequent biosynthetic steps, which correspond to modifications of the aromatic ring, are carried out by a multi-protein complex (named complex Q), associated with the mitochondrial inner membrane on the matrix side ^3^. The exact sequence of reactions that modify the aromatic ring is still a matter of debate. Traditional views, based on yeast data, suggested that the first reactions consist of the hydroxylation of position C5 of nP-HB catalysed by COQ6, and the subsequent methylation catalysed by COQ3, followed by the C1 decarboxylation and hydroxylation steps ^6,7^ (Fig. 1A). However, this sequence of reactions may be flexible since we reported that human and yeast cells with deficient COQ6 accumulate 3-polyprenyl-1,4-benzoquinone (PBQn) indicating that, at least in some situations, the C1 decarboxylation and C1 hydroxylation are the first modifications of the ring to take place ^7,8^ (Fig 1). Irrespective of the sequence of reactions, the enzymes that catalyze the C1 decarboxylation and C1 hydroxylation in eukaryotes are still uncharacterized ^3,9^. In bacteria such as *E. coli*, these reactions are carried out by two distinct enzymes, UbiD (a decarboxylase that uses a prenyl-FMN cofactor produced by UbiX ^10^) and UbiH (a hydroxylase) ^11^, which have no eukaryotic counterparts. Proteomic analyses of complex Q in eukaryotic cells failed to identify candidate proteins that could catalyze these reactions ^6,12^.

**Figure 1.**
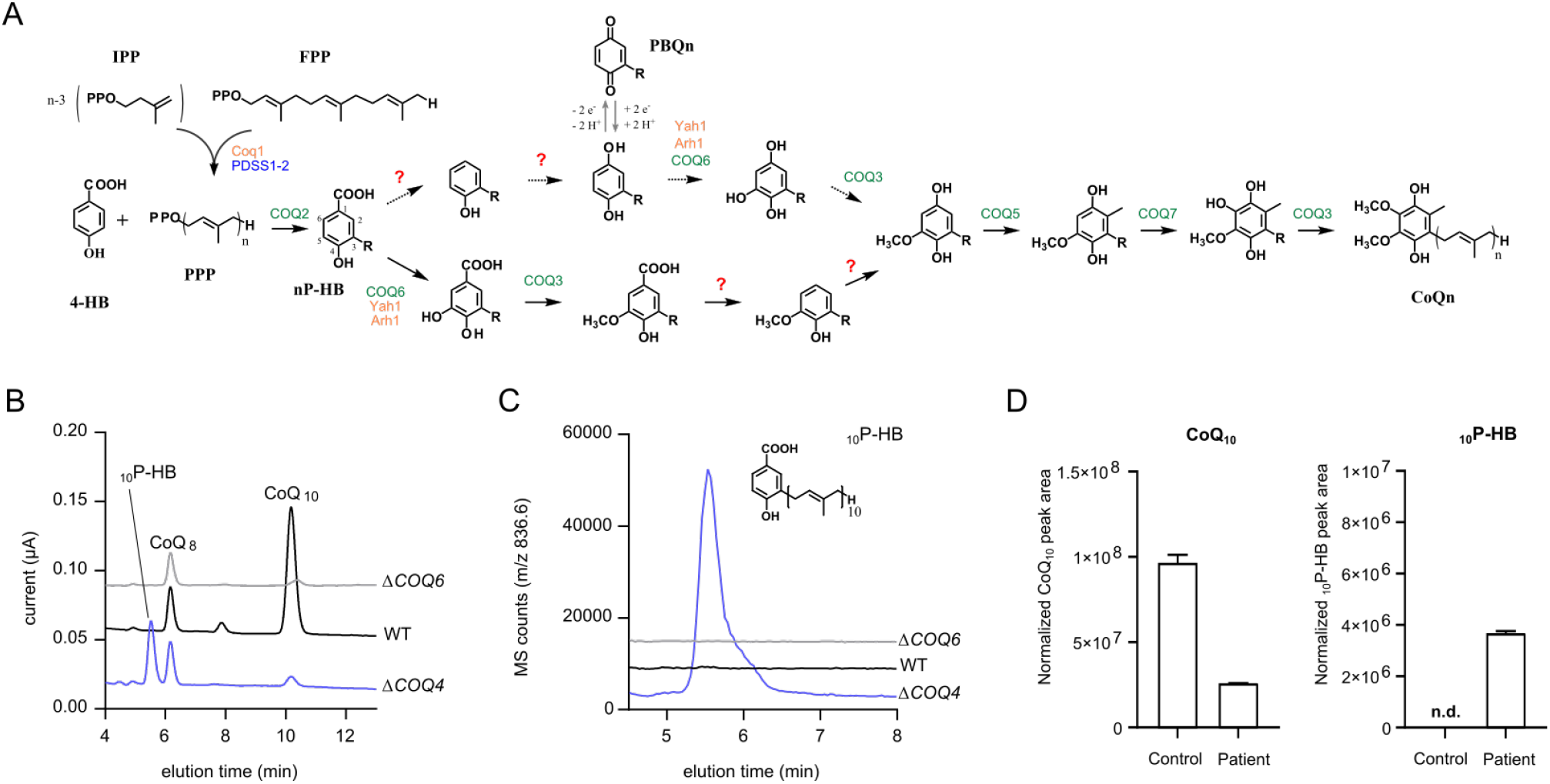
COQ4-deficient human cells accumulate _10_P-HB. A) The biosynthetic pathway of CoQ in eukaryotes. Proteins shared by *S. cerevisiae* and humans are in green, and those specific to *S. cerevisiae* or humans are in orange and blue, respectively. 4-hydroxybenzoic acid (4-HB) is the aromatic precursor of CoQ and the numbering of the carbon atoms used throughout this study is shown for polyprenyl-4-HB (nP-HB). Isopentenyl pyrophosphate (IPP) and farnesyl pyrophosphate (FPP) are building blocks for the synthesis of the polyprenyl pyrophosphate tail (PPP) which is added onto 4-HB by Coq2. The polyprenyl chain (n=6 for *S. cerevisiae*, n=10 for humans) is depicted by R on all intermediates derived from nP-HB. The uncharacterized steps for C1-decarboxylation and C1-hydroxylation are indicated with red question marks. An alternative order for the early steps is depicted by dashed arrows. The biosynthetic intermediates are represented in reduced form but they may also exist in their oxidized form as depicted for polyprenyl-1,4-benzoquinone (PBQn). B-C) Superimposition of electrochomatograms (B) and chromatograms of single ion monitoring (m/z 836.6 in positive mode) of _10_P-HB in lipid extracts from WT, Δ*COQ4* and Δ*COQ6* HEK293 cells (0.2 mg protein), with CoQ_8_ used as standard. D) Targeted mass spectrometry quantification of CoQ_10_ (left) and _10_P-HB from human primary fibroblast samples normalized to a CoQ_8_ internal standard. The data are representative of biological triplicates (B-D).

In addition to unassigned catalytic steps, gaps in our knowledge of eukaryotic CoQ biosynthesis also relate to the poorly characterized function of some of complex Q proteins, such as COQ4. The *COQ4* gene is relevant for human disease because its mutations are one of the most common causes of primary coenzyme Q deficiency ^13^. They are associated with a variety of clinical pictures ranging from fatal neonatal multisystem disorders ^14^, to childhood-onset cerebellar ataxia ^15^, or isolated retinitis pigmentosa ^16^. Interestingly, *COQ4* patients usually display a lower response to oral CoQ_10_ supplementation compared to patients with other COQ gene defects ^17^, hence the importance of understanding the precise function of COQ4.

COQ4 homologues exist in virtually all eukaryotes and potentially in some bacterial species, but not in *E. coli* ^18^. Based on the demonstration that *S. cerevisiae coq4* mutants failed to assemble complex Q and did not synthesize CoQ, COQ4 was proposed to fulfill a structural role in the organization of complex Q ^18^. However, the biosynthetic complex is destabilized in most yeast strains deleted for *coq* genes, and CoQ, or at least late biosynthetic intermediates like demethoxy-CoQ, seem important for complex Q stability ^19^. COQ4 proteins display a conserved motif, HDxxH(x)_11_E, which has been proposed to be a zinc-binding domain ^18^. Consistent with this hypothesis, the aspartic acid and glutamic acid residues of the HDxxH(x)_11_E motif coordinate two magnesium ions in the crystal structure of a cyanobacterial homologue of COQ4, Alr8543 from *Nostoc sp. PCC 7120* (PDB: 6E12) (Figure S1A), which has been used in the past to model eukaryotic COQ4 ^20^. Interestingly, mutation of the same glutamic acid residue in the *coq4-1* yeast strain impaired the function of COQ4 without affecting its stability ^18^. Adjacent to the metal coordinating pocket, the 6E12 crystal structure shows a hydrophobic tunnel with a bound molecule in which an oleic acid aliphatic chain is bound (Fig. S1B). This tunnel seems suitable to house the polyisoprenoid side chain of CoQ, potentially positioning the aromatic ring in close proximity to the metal-binding site. Based on these elements, we hypothesized that the function of COQ4 might not be limited to a structural role. Here, we provide evidence that COQ4 acts as an oxidative decarboxylase, substituting in a single step the carboxylic acid group with a hydroxyl group on carbon 1 of CoQ precursors.

## RESULTS

### Cells lacking COQ4 accumulate 3-decaprenyl-4-hydroxybenzoic acid, the carboxylated precursor of CoQ_10_

We have previously generated a HEK293 cell line in which the *COQ4* gene was disrupted by TALEN nucleases (Δ*COQ4* cells). These cells do not produce any CoQ_10_, but they incorporate some exogenous CoQ_10_ from the culture medium ^21^. We analysed lipid extracts from WT and Δ*COQ4* cells by HPLC. Electrochemical detection (ECD) revealed a strong decrease of CoQ_10_ in Δ*COQ4* cells and the presence of a compound that eluted at 5.6 min, right before the CoQ_8_ standard (Fig 1B). This compound was not detected in WT or Δ*COQ6* cells and was characterized by HPLC-coupled mass spectrometry. Mass scanning (m/z 660-900) in positive mode revealed a prominent ion at m/z 836.7 at a retention time of 5.6 min in the extracts of Δ*COQ4* cells (Fig S2A). A signal at 5.6 min was indeed specifically detected in Δ*COQ4* cells with single ion monitoring for m/z 836.6 (Fig 1C), a mass (M + NH_4_^+^) compatible with that of 3-decaprenyl-4-hydroxybenzoic acid (_10_P-HB). The compound also showed a MS signal at m/z 817.6 in negative mode, consistent with the ionization of the carboxyl moiety by loss of a proton (M-)(Fig S2B). Overall, these data show that Δ*COQ4* cells are impaired for the biosynthesis of CoQ_10_ and accumulate _10_P-HB, the product of the reaction catalysed by COQ2. A compound with a mass compatible with _10_P-HB was found also in cultured skin fibroblasts from a patient with primary CoQ deficiency due to COQ4 mutations ^15^ and was absent in control cells (Fig 1D). These observations can have two explanations. The absence of COQ4 might disrupt the integrity of complex Q, preventing the occurrence of all subsequent ring modifications, leading to an indirect accumulation of the mentioned precursor. Alternatively, we considered the hypothesis that COQ4 could have a direct role in CoQ biosynthesis, catalysing the decarboxylation of the C1 carbon of the aromatic ring.

### COQ4 binds zinc and metal binding is essential for function

The human COQ4 protein (hCOQ4) lacking the mitochondrial targeting sequence ^22^ and containing an N-terminal His Tag was expressed in *E. coli*. Although hCOQ4 was mainly found in inclusion bodies, we were able to partially solubilize it using 1% sarcosyl in the elution buffer (Fig S2C-D). Atomic absorption experiments showed that purified recombinant hCOQ4 treated with a zinc chloride solution retained approximately two zinc ions per protein (Fig 2A). We then mutagenized the aspartate in position 164 within the HDxxH(x)_11_E putative Zinc-binding domain (D164A). The D164A mutant was unable to complement respiratory growth in Δ*COQ4* yeast (Fig S2E) and the protein was virtually unable to bind zinc (Fig 2A). Interestingly, apo and holo hCOQ4 displayed far UV circular dichroism spectra comparable to those of the D164A mutant (Fig 2B), suggesting that the purified proteins maintained their fold irrespective of zinc binding.

**Figure 2.**
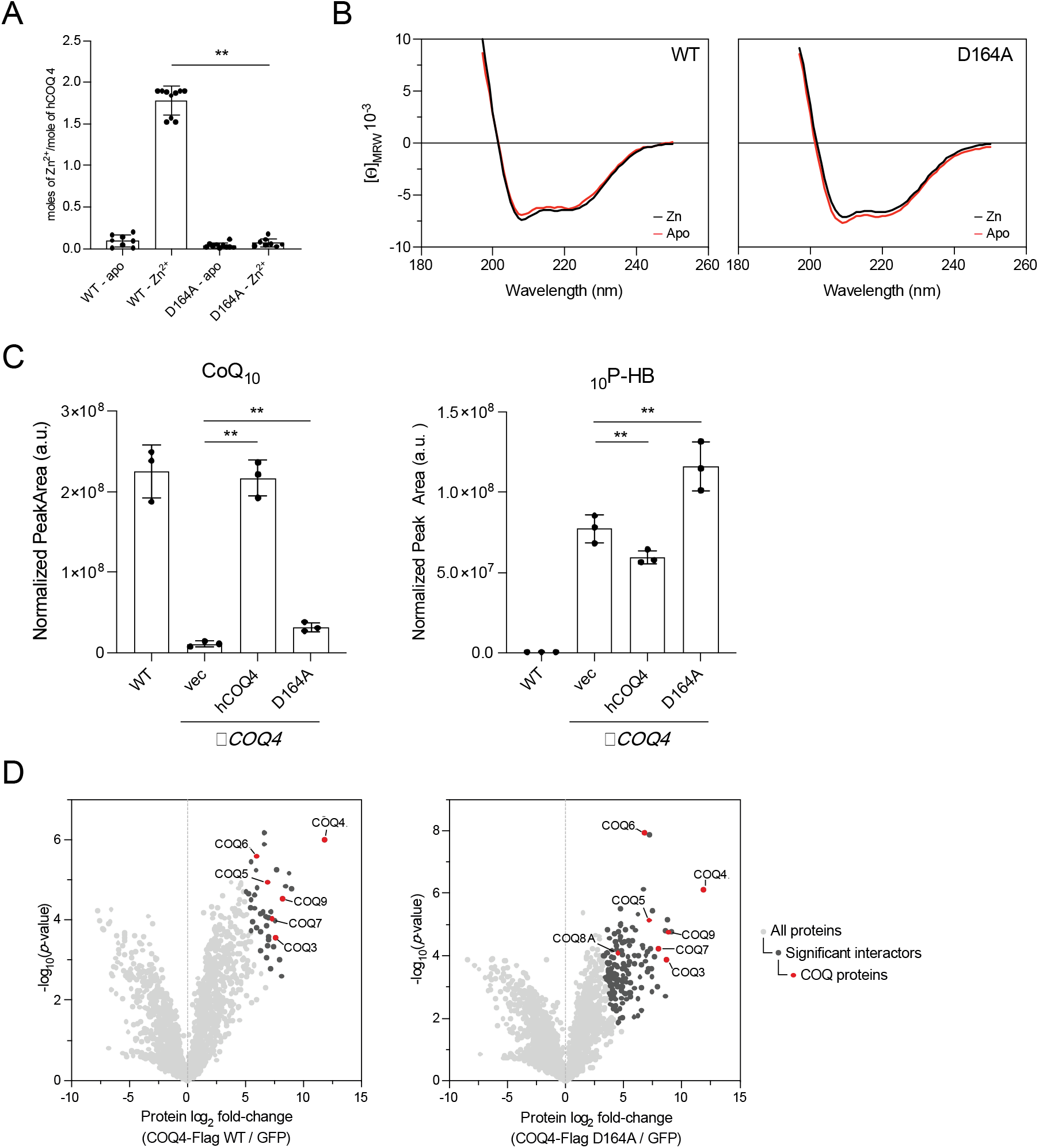
The D164A mutation impairs COQ4 function and metal binding, but does not compromise stability or interaction with complex Q. A) Zn^2+^/protein molar ratio of wild type and mutant COQ4. Proteins (3 µM) were treated with 10 mM EDTA (Apo) or with 100 µM ZnCl_2_ (Zn), followed by extensive dialysis against 25 mM Tris-HCl, pH 8.0, 100 mM NaCl. Zinc ion concentration was determined by atomic absorption spectrometry. **\*\*=**p<0.05. B) Far UV CD spectra of human recombinant COQ4 wild type and of the D164A mutant. Proteins (0,4 mg/ml) were treated with 10 mM EDTA (Apo) or with 100 µM ZnCl_2_ (Zn), followed by extensive dialysis against 25 mM Tris-HCl, pH 8.0, 100 mM NaCl. C) Targeted LC-MS-MS measurements of CoQ_10_ and _10_P-HB in WT or Δ*COQ4* HEK293 expressing a GFP control empty vector, hCOQ4 WT, or hCOQ4 D164A, shown as mean ± SD (n=3). **p < 0.05, two-sided Student’s *t*-test. D) Affinity enrichment mass spectrometry-based interaction profile of hCOQ4-Flag WT (left) and hCOQ4-Flag D164A (right) versus GFP control in WT HEK293 cells. Results representative of biological triplicate. Significant interactors, as determined by a two sample t-test with FDR < 0.05, are shown in dark gray, and COQ proteins are highlighted in red.

To test the functional importance of the zinc-binding domain, hCOQ4 WT or D164A was expressed in Δ*COQ4* HEK293 cells and CoQ_10_ levels were analysed by targeted mass spectrometry. Expression of hCOQ4 WT was sufficient to fully rescue the CoQ_10_ defect of Δ*COQ4* cells, however hCOQ4 D164A expression only marginally increased CoQ_10_ levels (Fig 2C). Consistent with previous results, Δ*COQ4* cells with empty vector accumulate _10_P-HB, whose levels are slightly decreased upon expression of hCOQ4 WT, though this may have been hindered by poor transient transfection efficiency of the constructs (Fig 2C). While a substantial increase in CoQ_10_ levels was observed upon expressing hCOQ4 WT but not D164A in Δ*COQ4* cells, a similar trend in respiratory rate was observed by Seahorse analysis, with the modest rescue again stemming from the limitations of the low transfection efficiency (Fig S2F). To interrogate if the D164A mutation disrupted the previously described scaffolding role of COQ4 in complex Q ^23^, we performed affinity enrichment mass spectrometry (AE-MS) of hCOQ4-FLAG WT and hCOQ4-FLAG D164A in WT HEK293 cells. Both WT and D164A COQ4 captured the same interaction profile of proteins involved in CoQ biosynthesis (Fig 2D), consistent with previous AE-MS experiments describing the nature of complex Q ^12^. Taken together, these experiments demonstrate that mutating the zinc binding domain of COQ4 disrupts its catalytic activity while maintaining its ability to participate structurally in complex Q.

### COQ4 rescues defects in C1-decarboxylation and C1-hydroxylation steps in the E. coli CoQ_8_ biosynthetic pathway

To verify if COQ4 might be involved in the decarboxylation step of the eukaryotic CoQ biosynthesis pathway, we set up a heterologous complementation assay with *E. coli* cells lacking the *ubiD* gene that encodes the decarboxylase of the bacterial CoQ pathway. *E. coli* Δ*ubiD* cells are deficient in CoQ_8_ ^24^ and consequently present a growth defect on respiratory medium but not on fermentation medium. Heterologous expression of both yeast COQ4 (yCOQ4) and hCOQ4 increased the respiratory growth of Δ*ubiD* cells, whereas the functionally inactive D164A mutant did not (Fig 3A). The respiratory growth reflected a substantial increase in CoQ_8_ levels, to 14 and 18% of WT levels for hCOQ4 and yCOQ4 (Fig 3B). This increase was found to be specific to COQ4 because expression of other COQ biosynthetic proteins from yeast (COQ3, COQ5, COQ6, and COQ9) induced no significant rescue of CoQ_8_ levels (Fig 3C). These data establish that COQ4 is sufficient to complement the defective decarboxylation step in the CoQ pathway of *E. coli* Δ*ubiD* cells.

**Figure 3.**
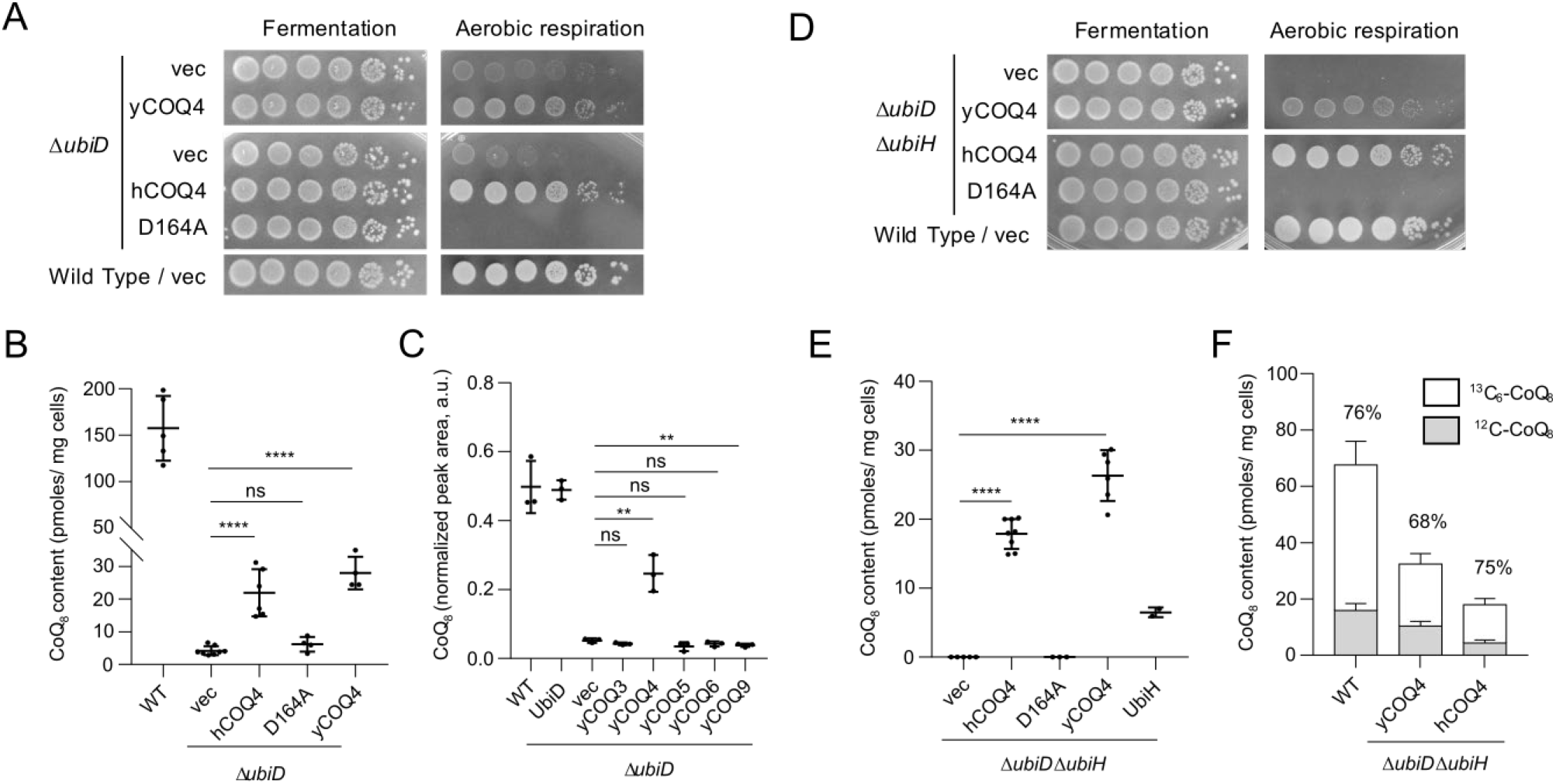
COQ4 complements defects in C1-decarboxylation and C1-hydroxylation in *E. coli*. A,D) Plate-growth assay of serial dilutions of Δ*ubiD* (A) or Δ*ubiD* Δ*ubiH* (D) cells containing empty vectors (vec) or vectors expressing yeast Coq4 (yCOQ4), human COQ4 (hCOQ4) or the human COQ4 D164A allele (D164A). B,E) CoQ_8_ quantification by HPLC-ECD-MS of Δ*ubiD* (B) or Δ*ubiD* Δ*ubiH* (E) cells containing vectors described in A, mean ± SD. **\*\*\*\*=**p<0.0001, ns = non-significant by unpaired t-test. CoQ_8_ levels for WT cells are similar in both experiments. C) CoQ_8_ quantification by targeted LC-MS-MS of Δ*ubiD* cells containing vectors expressing yCOQ4 or other *S. cerevisiae* CoQ biosynthetic proteins. **=p<0.05, ns = non-significant. F) Labelling of CoQ_8_ (percentage above the bars) from the ^13^C_7_-4HB precursor, mean ± SD (n = 3 to 4).

Since the C1-hydroxylase of the eukaryotic CoQ pathway remains uncharacterized (Fig 1), we tested whether this function might also be carried out by COQ4. *E. coli* cells lacking the C1-hydroxylase UbiH are deficient in CoQ_8_ and accumulate the biosynthetic intermediate octaprenyl-phenol with a decarboxylated C1 ^25^. Neither yCOQ4 nor hCOQ4 complemented the respiratory growth defect or the CoQ_8_ levels of Δ*ubiH* cells (Fig S3A-B). However, both genes complemented the phenotypes of *E. coli* cells lacking both the *ubiD* and the *ubiH* genes (Δ*ubiD* Δ*ubiH*), and again the nonfunctional hCOQ4-D164A mutant was incapable of doing so (Fig 3D-E).

We verified that CoQ_8_ synthesis in Δ*ubiD* Δ*ubiH* cells expressing COQ4 occurred from the normal 4-HB precursor. When ^13^C_7_-4HB was added to the growth medium, we observed a comparable labelling of CoQ_8_ in WT cells and in Δ*ubiD* Δ*ubiH* cells expressing COQ4 (Fig 3F). Thus, we conclude that cells expressing COQ4 use 4-HB as a precursor of CoQ_8_, which supports that the C1-decarboxylation and the C1-hydroxylation are taking place. Finally, we found that the activity of yCOQ4 and hCOQ4 depends on O_2_ since these proteins did not increase CoQ_8_ levels in Δ*ubiD* or Δ*ubiD* Δ*ubiH* cells grown under anaerobic conditions (Fig S3C). Overall, our data support a model in which COQ4 is able to perform a coupled decarboxylation and hydroxylation of position C1 in the CoQ biosynthesis pathway. We note that the presence of a carboxyl group on C1 seems necessary to the activity of COQ4 as C1 was not hydroxylated if it had already been decarboxylated by UbiD, as is the case in Δ*ubiH* cells.

### COQ4 synthesizes polyprenyl-1,4-benzoquinone in Corynebacterium glutamicum

*C. glutamicum*, an actinobacterium of biotechnological interest, synthesises a single isoprenoid quinone, MK_n_(H_2_), a partially saturated form of menaquinone ^26^ (Fig 4A). Recently, extensive genetic modification of *C. glutamicum* coupled to expression of the *ubi* genes from the *E. coli* CoQ biosynthesis pathway allowed production of CoQ_10_ in strain 413 ^27^. We reasoned that *C. glutamicum* would be the organism of choice to test the activity of COQ4, given the absence of endogenous CoQ biosynthesis genes. In comparison to the control strain 401, strain 405 expressing a decaprenyl synthase DdsA (a COQ1 homolog from *Paracoccus denitrificans*) and *E. coli* UbiA (the homolog of COQ2) produced several additional compounds that fall into two categories: polyprenyl-4-hydroxybenzoic acids (_8-11_P-HB) eluting between 3 and 7 min, and MK_8-11_(H_2_) eluting between 10 and 25 min (Fig S3D) ^28^. For both compounds, 9 and 10 were the most abundant isoprenologs (Fig S3D). Expression of yCOQ4 in strain 405 yielded several new peaks between 6 and 13 min that corresponded to isoprenologs of polyprenyl-1,4-benzoquinone (PBQ_8-11_, Fig 4B), as determined by mass spectrometry (Fig 4C and data not shown). This result established that yCOQ4 decarboxylated and hydroxylated position C1 of the _8-11_P-HB substrates. Expression of the *E. coli* decarboxylase UbiD (together with UbiX, which synthesizes the prenyl-FMN cofactor of UbiD ^10^) in strain 405 caused the complete conversion of _9-10_P-HB (Fig 4D) into nonaprenyl- and decaprenyl-phenol (_9_PP and _10_PP, the C1-decarboxylated products) (Fig 4E) without formation of any PBQ_9-10_ (Fig 4C) (the C1-hydroxylated product). Interestingly, _9_PP or _10_PP were not detected in the strain expressing yCOQ4 (Fig 4E), strongly suggesting that the conversion of _9-10_P-HB into PBQ_9-10_ occurred in a single step, via an oxidative decarboxylation. Together, these results demonstrate that COQ4 decarboxylates nP-HB into a product with an oxygenated substituent at C1, either a hydroxyl or a ketone (PBQn) depending on the redox state of the molecule (Fig 4A).

**Figure 4.**
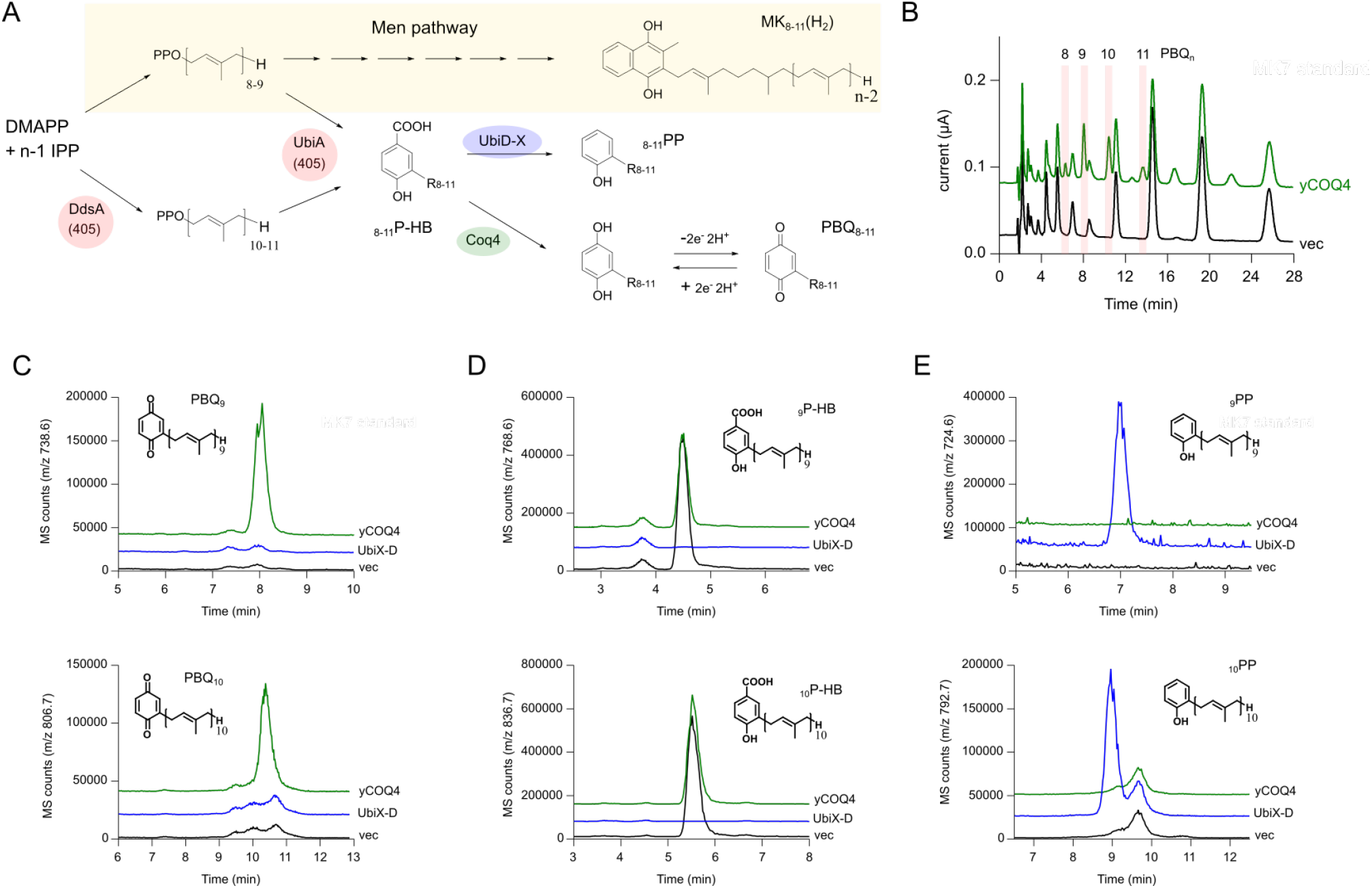
COQ4 synthesizes polyprenyl-1,4-benzoquinone in *Corynebacterium glutamicum*. A) Endogenous MK_n_(H_2_) biosynthesis pathway in *C. glutamicum* (highlighted in beige) and artificial polyprenyl compounds obtained via expression of heterologous genes. Strain 405 expresses *ubiA* and *ddsA*, and produces _8-11_P-HB. Additional expression of *E. coli ubiD* and *ubiX* or *S. cerevisiae* COQ4 (yCOQ4) yields, _8-11_PP or PBQ_8-11_, respectively. B) Electrochromatograms obtained by HPLC-ECD-MS analysis of lipid extracts from strain 405 containing the empty plasmid pEC-XT99A (vec) or the pEC-XT99A-yCOQ4 plasmid. The polyprenyl-1,4-benzoquinones (PBQ_8-11_) synthesized specifically in the presence of yCOQ4 are labelled based on their identification by MS (see fig 4C). C-E) Single ion monitoring analyses of nonaprenyl and decaprenyl forms of PBQ_n_ (C), nP-HB (D), and nPP (E) in lipid extracts of strain 405 expressing yCOQ4 (pEC-XT99A-yCOQ4), UbiX and UbiD (pEC-XT99A-ubiDX), or a control plasmid (pEC-XT99A, vec). Results representative of biological triplicates (B-E).

## DISCUSSION

The exact function of COQ4 within CoQ biosynthesis and the nature of the enzymes responsible for the C1 decarboxylase and hydroxylase activities in the eukaryotic CoQ pathway have remained elusive for many years. We now provide evidence that COQ4 carries out these two reactions in single step as an oxidative decarboxylation reaction. In fact, we showed that expression of *COQ4* complemented *E. coli* strains deficient either for the decarboxylase (Δ*ubiD*) or for the decarboxylase and the C1-hydroxylase (Δ*ubiD* Δ*ubiH*). These data suggest that COQ4 converts octaprenyl-4-hydroxybenzoic acid (_8_P-HB, the C1-carboxylated precursor), which accumulates in Δ*ubiD* and Δ*ubiD*Δ*ubiH* strains, into a molecule with an oxygen-substituted C1 position like octaprenyl-1,4-benzoquinone (PBQ_8_) or its reduced form (see Fig 1A). Since COQ4 is ineffective in the *E. coli* Δ*ubiH* strain that accumulates octaprenyl-phenol (_8_PP), we suggest that COQ4 requires a carboxyl group on C1 and does not function as a *bona fide* C1 hydroxylase, but rather as an oxidative decarboxylase, similar to the VibMO1 protein from *Boreostereum vibrans*, a protein which belongs to the family of FAD-dependent monooxygenases ^29^. Experiments in *C. glutamicum* support this view since cells expressing COQ4 synthesized PBQ_9-10_ without any accumulation of _9/10_PP, contrary to cells expressing UbiD-X in which it was the only product (Fig 4). In addition, a putative requirement of other COQ proteins to assist the oxidative decarboxylation of C1 is ruled out because COQ4 alone was sufficient to form PBQ_8-11_ from _8-11_P-HB in *C. glutamicum*, an organism that does not produce CoQ naturally and therefore does not contain endogenous COQ proteins.

The fact that Δ*COQ4* cells accumulate mainly _10_P-HB (the product of COQ2) and that Δ*COQ6* cells accumulate PBQ_10_ (the product of COQ4) support the hypothesis that, at least in our HEK293 cell model, the oxidative decarboxylation by COQ4 occur preferentially before the C5 hydroxylation by COQ6 ^8^. We therefore propose an updated eukaryotic CoQ biosynthesis pathway with this sequence of reactions (path 1, Fig S4). It should be noted that the sensitivity of our HPLC-mass spectrometry analyses is limited, therefore we cannot exclude the formation of small amounts of other biosynthetic intermediates, which may result from alternative sequences of reactions in the knockout cells. Compounds that accumulate in cells deficient for *COQ* genes may not necessarily correspond to the substrates of the deficient COQ enzymes as some reactions may occur “out of order” when the CoQ biosynthesis pathway is impaired. This is exemplified by DMQ_10_, the substrate of COQ7 and penultimate intermediate of the CoQ pathway, which was detected in low amounts in COQ3-COQ6 siRNA cells, but which accumulated at much higher levels in the COQ7 siRNA cells ^30^. We consider that the “traditional” order of the CoQ pathway (path 2, Fig S4) is also compatible with our findings since previous studies in yeast showed that vanillic acid can replace 4-HB as a substrate of CoQ ^7^. As vanillic acid is prenylated by COQ2 before being decarboxylated on C1 ^31^, yCOQ4 is necessarily able to process a substrate with a methoxyl group on C5 (path 2 in Fig S4). As such, COQ4 promiscuity may allow various sequences of reactions to take place. Interestingly, the order of the reactions in the bacterial CoQ pathway was recently proposed to differ between *E. coli* and *Rhodobacter capsulatus*, in which a new flavin-dependent hydroxylase was shown to function as a C1-hydroxylase and also as a C1-decarboxylase when UbiD was inactivated ^32^.

A number of open issues still exist around COQ4. First, the exact mechanism of the oxidative decarboxylation reaction is not clear. Molecular oxygen is required as demonstrated by the lack of complementation of *E. coli* mutants in anaerobic conditions, and by labelling experiments in yeast, which showed that all three oxygen atoms added on CoQ precursors are derived from O_2_ ^33^. Furthermore, we still do not know which metal ion is bound by COQ4 under physiological conditions. Alr8543 was crystallized with magnesium ions, and we showed zinc binding in recombinant human COQ4. In fact, the geometry of the binuclear metallic site of COQ4 is similar to that of binuclear metallohydrolases, which are known to bind different types of metal ions (Fe, Zn, Mn, Co) in their active site ^34^. Therefore, the COQ4 active site may accommodate various metal ions, but finding those with physiological relevance will likely necessitate the development of an *in vitro* activity assay. Our repeated attempts to detect *in vitro* activity with purified recombinant COQ4 using different combination of metals (zinc, iron, copper), cofactors (NADH, NADPH, FAD, ATP), and substrates (4-HB, vanillic acid, polyprenyl vanillic acid derived from yeast lipid extracts) were unsuccessful. The presence of a hydrophobic tunnel next to the putative active site of the cyanobacterial homolog of COQ4 suggests that a polyprenyl side chain on the substrate might be necessary for the reaction to take place. However, the hydrophobicity of polyprenylated substrates complicates their use in *in vitro* assays with aqueous buffers. Moreover, since COQ4 is associated with the inner mitochondrial membrane ^23^, the presence of lipids could also be critical for enzymatic activity. Several of the issues discussed above might be clarified if structural data were available, which is unfortunately not the case for any eukaryotic COQ4 homologs.

Even though we have obtained only genetic data in this study, our demonstration of COQ4 activity in two different *in vivo* models provides strong support for our conclusions. In particular, the activity observed in the non-natural CoQ producer *C. glutamicum* is undoubtedly linked to the heterologously expressed yCOQ4. Our conclusion that COQ4 carries out, in a single step, the oxidative decarboxylation of the C1 carbon of CoQ precursors fills a major gap in the eukaryotic CoQ biosynthesis pathway since specific enzymes have now been assigned to all the steps required for ring modification.

## ACKNOWLEDGEMENTS

This work was supported by grants from the ANR to FP (ANR-19-CE44-0014O2-TABOO), from the NIH to DJP (R35GM131795), from Telethon Italy (GGP13222), from Fondazione IRP Città della speranza (to LS), from Fondazione CARIPARO (20/19 FCR) to ET, and from European Regional Development Fund (FEDER)/Andalusian Regional Government (FEDER operational program 2014– 2020; P18-RT-4572).

## AUTHORS CONTRIBUTION SECTION

Conceptualization, L.S., F.P.; methodology, L.S., F.P., P.N., G.Z., R.G., F.T., M.A.D., C.A., G.B.C., E.T., D.J.P., W.F.W.; investigation, L.P., L. M., A.B., A.K.B., R.G., K.K.F., B.R., M.R.M.; formal analysis, F.T., A.K.B., D.J.P., G.B.C., W.F.W., L.S., F.P.; resources, E.T., L.S., F.P., W.F.W.; writing – original draft, L.S., F.P.; writing – review & editing, F.P., D.J.P., F.T, W.F.W., C.A., G.B.C., R.G.; supervision, G.B.C., M.A.D., F.T.; funding acquisition, F.P., D.J.P, L.S., E.T.

## STAR METHODS

### Resource availability

#### Lead contact

Further information and requests for resources and reagents should be directed to and will be fulfilled by the L.S.

#### Materials availability

All unique/stable reagents generated in this study are available from the lead contact upon reasonable request.

#### Data and code availability

All data were deposited in Cell-Press-recommended, datatype-specific repositories and made publicly accessible.

- The raw proteomic data generated during the current study have been deposited at MassIVE repository and are publicly available under accession code MSV000093201 as of the date of publication. Accession numbers are also listed in the key resources table.

This paper does not report original code.

Any additional information required to reanalyze the data reported in this paper is available from the lead contact upon request.

### Experimental model and subject details

#### Cell lines

##### Human Cells and Culture conditions

Δ*COQ4*, Δ*COQ6* cell lines, and patients fibroblasts have been reported previously ^8,15,35^. Fibroblasts were cultured in DMEM High Glucose (Gibco™ 11965092), supplemented with 10% FBS, and HEK293 cells were cultured in DMEM High Glucose (Gibco™ 11965092), supplemented with 10% FBS, 1% penicillin-streptomycin, 1 mM pyruvate, 2 mM glutamine, and 10 µM uridine.

##### Microbes

###### E. coli

*E. coli* strains are listed in the Key resources table. Unless otherwise stated, *E. coli* cultures were performed in lysogeny broth medium (LB) at 37°C with 180 rpm shaking. Anaerobic cultures were performed in Hungate tubes containing 12 mL LB medium (with 0.5 mL/L antifoam B emulsion, Sigma) deoxygenated by argon (O_2_<0.1 ppm) bubbling for 25 min before autoclaving. Hungate tubes were inoculated through the septum with 100 µL of overnight precultures taken with disposable syringes and needles from closed Eppendorf tubes filled to the top. Ampicillin (50 mg/L) or kanamycin (25 mg/L) were added from stock solutions (1,000X, sterilized through 0.22 µm filters and stored at −20°C) to maintain plasmid selection. Media contained 100 μM IPTG or 0.05% arabinose for induction of the pTrc-yCOQ4 or pBAD-hCOQ4 plasmids, respectively.

###### S. cerevisiae

*S. cerevisiae* lacking *COQ4* (*W303Δcoq4)* was transformed using the PEG-lithium acetate method as previously described ^36^. Transformants were selected and maintained in minimum medium (SM Glu) containing 2% glucose, 0.17% yeast nitrogen base without amino acids, 0.5% ammonium sulfate, leucine 60 mg/L, tryptophan 20 mg/L.

###### C. glutamicum

*C. glutamicum* strains 401 and 405 were obtained from strain UBI400 ^28^ by transforming and empty pRG_Duet2 or pRG_Duet2-*ddsA*_Pd_-*ubiA*_Ec_, respectively. Transformation was carried out via electroporation ^37^ at 2.5 kV, 200 Ω, and 25 µF. Strain 405 was additionally transformed with plasmids pEC-XT99A, pEC-XT99A-yCOQ4 or pEC-XT99A-ubiDX.

### Method details

#### Plasmids construction

Oligonucleotides used in this study are listed in Table 1. Human *COQ4* devoid of the mitochondrial targeting sequence (hCOQ4) was amplified from pCRII-TOPO-COQ4 (which contains the entire coding region, the STOP codon and 10 nucleotides of the 3’ UTR) ^22^ using primers hCOQ4_HisT_NdeI_For (which encodes the His Tag and enterokinase consensus) and hCOQ4_HindIII_R, digested with NdeI and HindIII (NEB) and cloned into the pET21a vector (Snapgene) cut similarly. The D164A mutant was obtained by site directed mutagenesis using the QuikChange lightning site-directed mutagenesis kit (Agilent) which employs a high-fidelity Pfu Ultra DNA polymerase and primers D164A_F and D164A_R. The PCR products were digested with DpnI (Agilent) and then used to transform *E. coli* Mach1 competent cells. For subsequent cloning into pBAD24, *COQ4* was amplified from pET21a-hCOQ4 or pET21a-hCOQ4_D164A using primers hCOQ4_BAC_ NheI_F and hCOQ4_HindIII_ R, digested with NheI and HindIII, and cloned into the pBAD vector similarly digested. The resulting pBAD-hCOQ4 plasmid contained hCOQ4 without the mitochondrial targeting sequence and without any tag. The open reading frame of the yeast COQ4 gene (*yCOQ4*) was amplified from the genomic DNA of the *S. cerevisiae* W303 strain using primers yCOQ4_Sac_Fw and yCOQ4_HIII_Rv. The PCR product was digested with SacI and HindIII and cloned into pTrc99a digested similarly to yield pTrc-yCOQ4. pTrc-UbiH was constructed by cloning the PCR-amplified *UBIH* gene (using primers UbiH5-EcoRI / UbiH3-HindIII and *E. coli* K-12 MG1655 genomic DNA as template) into pTrc99a after digestion with EcoRI and HindIII. All constructs were verified by Sanger sequencing. For yeast expression studies pCM189_COQ4 ^15^ was mutagenized using primers D164A_F and D164A_R as described above) to yield the D164A mutant (pCM189_COQ4^D164A^). Plasmids for other COQ biosynthetic proteins were prepared in the pET28b background with an ampicillin resistance cassette. Sequences for NΔ31 yCOQ3, NΔ30 yCOQ5, NΔ16 yCOQ9 were amplified from yeast vectors containing the full *S. cerevisiae* genomic coding regions for those genes. N-terminal truncations sites were determined using MitoFates ^38^. The PCR products for the COQ3/COQ5/COQ9 constructs were integrated into the empty pET28b using a HindIII and XhoI restriction enzyme (NEB) digest. The full-length COQ6 construct was amplified from a gBlock gene synthesis (IDT) containing a codon-optimized gene sequence for yeast COQ6 and was inserted into the pET28b backbone using Gibson cloning via the NEBuilder® HiFi DNA Assembly Cloning Kit (NEB). All four constructs were verified using Sanger sequencing.

*C. glutamicum* plasmids were constructed via Gibson Assembly ^28^ after gene amplification with Phusion High-Fidelity DNA polymerase (New England Biolabs, United Kingdom). The gene *ubiX* was amplified from genomic DNA of *E. coli* K-12 MG1655 using the primers ubiX-fw and ubiX-rv. The gene *COQ4* from *S. cerevisiae* was codon-harmonized ^39^, synthesized (Life Technologies GmbH, Darmstadt, Germany), and amplified using the primers yCOQ4-CG-fw and yCOQ4-CG-rv. The plasmid pEC-XT99A-*ubiD* ^27^ was cut using XbaI to construct pEC-XT99A-*ubiDX* and pEC-XT99A ^40^ was cut using BamHI to construct pEC-XT99A-yCOQ4.

Mammalian expression plasmids pcDNA3.1 GFP-FLAG and pcDNA3.1 hCOQ4-FLAG were generated and described previously ^12^. The pcDNA3.1 hCOQ4-FLAG construct was mutagenized using the QuikChange lightning site-directed mutagenesis kit (Agilent) and primers hCOQ4_D164A_F and hCOQ4_D164A_R to generate pcDNA3.1 hCOQ4-FLAG D164A.

#### Construction of *E. coli* strains

The chloramphenicol cassette was eliminated from the Δ*ubiD::cat* strain using plasmid pCP20 as described previously ^41^ to yield the Δ*ubiD* strain. The Δ*ubiD* Δ*ubiH* strain was constructed by introducing the *ubiH::kan* deletion into the Δ*ubiD* strain with P1 vir transduction ^42^ and selection for kanamycin resistance. The proper insertion of the cassette was verified by locus PCR with primers UbiHprom5 and UbiHterm3.

#### Mammalian cell culture and transfection

HEK293 WT and Δ*COQ4* cells were grown in Dulbecco’s modified Eagle’s medium (DMEM, LifeTechnologies) supplemented with 10% fetal bovine serum, 1% penicillin-streptomycin, 1 mM pyruvate, 2 mM glutamine, and 10 µM uridine. Cells were subcultured by trypsinization. For affinity enrichment mass spectrometry experiment, 7 x 10^6^ HEK293 cells were plated in a 15 cm dish and allowed to grow overnight. On day two, cells were transiently transfected with a mix of 20 µg plasmid (pcDNA3.1 GFP-FLAG, pcDNA3.1 hCOQ4-FLAG WT, or pcDNA3.1 hCOQ4-FLAG D164A), 72 µg linear polyethylenimine (PEI, PolySciences), and 900 µL Opti-MEM (LifeTechnologies). After 48 hours (Day 4), cells were washed with and harvested into phosphate-buffered saline (PBS), were collected at 2,000 rcf, were snap frozen in liquid nitrogen, and were stored at −80°C.

For CoQ_10_ measurements, 2.5 x 10^6^ HEK293 WT and Δ*COQ4* were plated in a 10 cm dish and allowed to grow overnight. On day two, cells were transiently transfected with a mix of 10 µg plasmid (pcDNA3.1 GFP-FLAG, pcDNA3.1 hCOQ4-FLAG WT, or pcDNA3.1 hCOQ4-FLAG D164A), 36 µg linear polyethylenimine (PEI, PolySciences), and 500 µL Opti-MEM (LifeTechnologies). After 48 hours (Day 4), cells were washed with and harvested into phosphate-buffered saline (PBS), were collected at 2,000 rcf, were snap frozen in liquid nitrogen, and were stored at −80°C.

#### Expression and purification of human COQ4

The expression of hCOQ4 and of the D164A allele from the pET21a plasmids was induced by adding 1mM isopropyl-1-thio-β-D-galactopyranoside to the culture of *E. coli* BL21 (DE3) cells. After incubation for 4 h at 30°C, cells were harvested by centrifugation (for 20 min, at 2400 g, at 4 °C), suspended in the lysis buffer (Na_2_HPO_4_ 20 mM, NaCl 0.5 M, imidazole 5 mM, DTT 1mM, pH 7.8), and lysed by sonication (Omni Sonic Ruptor 400 Ultrasonic Homogenizer, 30-second cycle: 5 seconds ON, 25 seconds OFF −> 14 cycles total). After centrifugation (for 30 min at 18000 g 4°C) and due to the low solubility of the protein, the inclusion bodies were solubilized O/N at room temperature in the lysis buffer added with 1% sarcosyl. After centrifugation (for 20 min, at 2400g, at 4 °C), the solubilized protein was isolated with a Ni affinity column (HisTrap HP, 5 mL, Cytiva) using an AKTA-FPLC and the eluate was dialyzed against a buffer containing NaCl 150 mM, Hepes 20 mM, DTT 1 mM, sarcosyl 0.1%, and ± ZnCl_2_ 100 µM. The excess of ZnCl_2_ was eliminated by extensive dialysis against the same buffer without zinc. The recombinant hCOQ4 was further purified with a gel filtration chromatography using a Superdez Yarra SEC-2000 column (Phenomenex). The sample purity and the molecular weight were checked by SDS/PAGE (see Fig S2C). Protein concentration was determined by UV absorption at 280 nm and by Brosted-Lowry method.

#### Circular dichroism spectroscopy

Measurements of optical ellipticity were made at room temperature on a Jasco J-710 spectropolarimeter in a 0.1 cm path-length quartz cell at a protein concentration of 0.4 mg/mL in 25 mM Tris-HCl, pH 8.0, 100 mM NaCl. Ellipticities were normalized to residue concentration using the relationship [θ]=(θobs/10)·(M/l*c) where θ obs is the observed ellipticity at a given wavelength, M is the mean residue mass, l is the cuvette path length in centimeters, and c is the protein concentration expressed as g/mL. All CD spectra resulted from an averaging of at least three scans and the final spectrum was corrected by subtracting the corresponding buffer spectrum obtained under identical conditions.

#### Analysis of zinc content

The zinc content of the hCOQ4 and hCOQ4-D164A proteins purified in the presence or absence of zinc was measured using a Perkin–Elmer 4000 atomic absorption flame spectrophotometer. Samples containing 3 µM of protein were measured after standardization in the linear range of 0–0.5 ppm of zinc and the buffer was used as blank. Analyses were performed in triplicate.

#### Yeast complementation assay

*S. cerevisiae W303Δcoq4* harboring the pRS423-yCOQ8 plasmid to overexpress yeast Coq8, was transformed with pCM189_COQ4 or pCM189_COQ4^D164A^. For plate dilution assays, each strain was cultured overnight in SM Glu medium. Optical densities (A_600 nm_) of harvested cells were adjusted to 0.2, and 2 μl of 1:10 serial dilutions were spotted onto agar plates (Bacto-agar 16 g/L) containing SM Glu, YPD (1% yeast extract, 2% peptone, 2% glucose) or YPGly (1% yeast extract, 2% peptone, 3% glycerol) media. The plates were imaged after incubation for 5 days at 30 °C.

#### *in vivo* COQ4 activity assays

##### *E. coli* serial dilution assay

Overnight cultures in LB medium with antibiotics for plasmid selection were adjusted to 1 OD_600_ and used for 10 fold serial dilutions. 5 μL drops were spotted on agar plates (Bacto-agar 16 g/L) with fermentation medium (LB + 0.2% glucose) or with respiration medium (synthetic medium^43^ supplemented with 0.4% succinate adjusted to pH 6). The plates contained either 100 µM IPTG or 0.05% arabinose for induction of pTrc-yCOQ4 or pBAD-hCOQ4, respectively. The plates were imaged after incubation for 20 hours at 37 °C in aerobic conditions (aerobic respiration) or inside a GasPak anaerobic jar fitted with a BD anaerobic sachet (fermentation).

##### COQ4 expression in *C. glutamicum*

Pre-cultures were cultivated in brain heart infusion (BHI) medium at 30 °C in baffled shake flasks at 120 rpm. Respective antibiotics were added in concentrations of 25 µg mL^-1^ kanamycin, 5 µg mL^-1^ tetracycline or 100 µg mL^-1^ spectinomycin. After approximately 16 hours, the cells were washed with TN buffer pH 6.3 (10 mM Tris-HCl, 150 mM NaCl), inoculated to an optical density (OD_600_) of 1 in CGXII minimal medium^37^ with 40 g L^-1^ glucose, 1 mM IPTG, and 0.25 µg mL^-1^ anhydrotetracycline (ATc), and cultivated for 72 hours at 30 °C and 120 rpm. 100 mg (wet weight) of cells were centrifuged at 20,200 *g* for 10 min and stored at −20 °C until quantification of CoQ biosynthetic intermediates.

#### Quantification of coenzyme Q and biosynthetic intermediates

##### Measurement by HPLC-ECD-MS

Non polar lipids were extracted from HEK293 cells, and CoQ_10_ and _10_P-HB were analyzed by HPLC-electrochemical detection-mass spectrometry as previously described ^8^ with the following modifications. For analyses in reducing conditions, a potential of −800 mV was used on the precolumn electrode. The cone voltage was set to −80 V for mass spectrometry detection in negative mode. For *E. coli* cells, extraction and analysis were as described previously ^44^.

##### Targeted Measurement by LC-MS/MS

For determination of the CoQ content in human primary fibroblasts and HEK293 cell rescue experiments, the frozen cell pellets were thawed and resuspended in 100 μL of PBS, then 5 uL of the resuspension was set aside for BCA assay to normalize the protein content. For *E. coli* pellets, the cell number was determined using optical density and the same number of cells was collected from each culture for centrifugation and freezing. After thawing, 50μL of 150mM KCl was added and used for resuspension. 600 μL of methanol with 0.1 μM CoQ_8_ (for human cell experiments) or 0.1 μM CoQ_6_ (for bacterial experiments) internal standard (Avanti Polar Lipids) was added to each sample. The cells were lysed by 5 min of shaking at 4 °C in a Disruptor Genie set to 3000 rpm. To extract lipids, 400 μL of petroleum ether was added to the sample and subjected to shaking as described previously for 3 min. Sample was spun 2 min. x 1000 g at 4°C and the top ether layer was transferred to a new tube. Extraction was repeated twice before the ether layers were combined and dried for 30 min under argon gas at room temperature. Extracted dried lipids were resuspended in mobile phase (78% methanol, 20% isopropanol, 2% 1 M ammonium acetate pH 4.4 in water) and transferred to amber glass vials with inserts. LC-MS analysis was performed using a Thermo Vanquish Horizon UHPLC system coupled to a Thermo Exploris 240 Orbitrap mass spectrometer. For LC separation, a Vanquish binary pump system (Thermo Fisher Scientific) was used with a Waters Acquity CSH C18 column (100 mm × 2.1 mm, 1.7 μm particle size) held at 35 °C under 300 μL/min flow rate. Mobile phase A consisted of 5 mM ammonium acetate in acetonitrile:H_2_O (70:30, v/v) with 125 μL/L acetic acid. Mobile phase B consisted of 5 mM ammonium acetate in isopropanol:acetonitrile (90:10, v/v) with the same additive. For each sample run, mobile phase B was initially held at 2% for 2 min and then increased to 30% over 3 min. Mobile phase B was further increased to 50% over 1 min and 85% over 14 min and then raised to 99% over 1 min and held for 4 min. The column was re-equilibrated for 5 min at 2% B before the next injection. Five microliters of the sample were injected by a Vanquish Split Sampler HT autosampler (Thermo Fisher Scientific), while the autosampler temperature was kept at 4 °C. The samples were ionized by a heated ESI source kept at a vaporizer temperature of 350 °C. Sheath gas was set to 50 units, auxiliary gas to 8 units, sweep gas to 1 unit, and the spray voltage was set to 3500 V for positive mode and 2500 V for negative mode. The inlet ion transfer tube temperature was kept at 325 °C with 70% RF lens. For targeted analysis, the MS was operated in positive parallel reaction monitoring mode with polarity switching acquiring scheduled, targeted scans to CoQ_10_ (m/z 863.6912), CoQ_8_ (m/z 727.566), CoQ_6_ (m/z 591.4408), and CoQ intermediates: DMQ_10_ (m/z 833.6806), DMeQ_10_ (m/z 849.6755), DDMQ_10_ (m/z 836.6915), _10_P-HB (m/z 817.6504) and BPQ_10_ (m/z 789.6555). MS acquisition parameters include resolution of 15,000, stepped HCD collision energy (25%, 30%, 40% for positive mode and 20%, 40%, 60% for negative mode), and 3s dynamic exclusion. Automatic gain control targets were set to standard mode. The resulting CoQ intermediate data were processed using TraceFinder 5.1 (Thermo Fisher Scientific).

##### C. glutamicum

CoQ biosynthetic intermediates were extracted from *C. glutamicum* cells (10-25 mg) and analyzed as described previously ^28^. Glass beads (100 μL), 50 μL of 150 mM KCl, and a volume of 2 mM MK_7_ solution (used as an internal standard, Sigma-Aldrich) proportional to the cell weight (2 μL/mg) were added to cell pellet. CoQ intermediates were extracted by adding 0.6 mL of methanol, vortexing for 10 min, then adding 0.4 mL of petroleum ether (boiling range 40–60 °C) and vortexing for 3 min. The phases were separated by centrifugation at room temperature for 1 min, 4,000 *g* and the upper petroleum ether layer was transferred to a fresh tube. The extraction was repeated with 0.4 mL petroleum ether, and petroleum ether layers were combined and dried under nitrogen. The dried samples were suspended in 100 μL ethanol and aliquots corresponding to 2 mg of cells wet weight were analyzed by reversed-phase HPLC with a C18 column (Betabasic-18, 5 μm, 4.6 × 150 mm ; Thermo Scientific) at a flow rate of 1 mL/min with a mobile phase composed of 25% isopropyl alcohol, 20% ethanol, 45% methanol and 10% of a mix of 89.9% isopropanol/10% ammonium acetate (1 M)/0.1% formic acid. A precolumn 5020 guard cell operated in oxidizing mode (E, +650 mV) and compounds were monitored by mass spectrometry with an MSQ Plus spectrometer and by electrochemical detection with an ESA Coulochem III detector equipped with a 5011A analytical cell (E1, −650 mV ; E2, +650 mV). The MSQ Plus was used in positive mode (probe temperature 400 °C, cone voltage 80 V) and single ion monitoring detected the following (NH_4_^+^ adducts, scan time 0.2 s) : _9_PP, m/z 724.1–725.1, 5–10 min ; _10_PP, m/z 792.2–793.2, 6–13 min ; _9_P-HB, m/z 768.1– 769.1, 2–7 min ; _10_P-HB, m/z 836.2–837.2, 3–8 min ; PBQ_9_, m/z 738.1–739.1, 5–10 min ; PBQ_10_, m/z 806.2–807.2, 6–13 min.

#### CoQ_8_ labelling in *E. coli* with ^13^C_7_-4HB

Overnight cultures in LB medium of MG1655 and Δ*ubiD* Δ*ubiH* cells containing pBAD-hCOQ4 or pTrc-yCOQ4 were used to inoculate (at OD_600_= 0.1) 10 mL LB cultures supplemented with 10 µM ^13^C_7_-4HB (Sigma) and the proper inductor. Cultures were stopped on ice when OD_600_ reached 0.9-1. Cells were collected and CoQ_8_ was extracted and analyzed by HPLC-MS ^43^ with single ion monitoring (6-10 min) for H^+^ adducts at m/z 727-728 (UQ_8_) and m/z 733-734 (^13^C_6_-UQ_8_).

#### Affinity Enrichment Mass Spectrometry

Pellets of HEK293 cells expressing WT or D164 versions of COQ4-Flag, or GFP-FLAG as a negative control, were lysed in 200 µL cold lysis buffer (20 mM HEPES, pH 7.4, 100 mM NaCl, 10% glycerol, 3% digitonin (Sigma), 1 mM DTT, and 1X cOmplete protease inhibitor cocktail (Roche). After periodic vortexing on ice, insoluble materials were pelleted (16,000 g, 10 min, 4°C) and the supernatant was retained. The protein concentration was quantified by BCA, and equal masses of cell supernatant were mixed with 30 µL pre-washed anti-FLAG magnetic beads for 2 h at 4°C with end-over-end agitation. Following incubation, beads were washed four times in wash buffer (20 mM HEPES, pH 7.4, 100 mM NaCl, 0.05% digitonin, 10% glycerol) and twice in final wash buffer (20 mM HEPES, pH 7.4, 100 mM NaCl) before being subjected to on-bead trypsin digest. The on-bead proteins were denatured with 2M urea in 200 mM Tris pH 8.0, then reduced with 5 mM DTT for 30 min at 56°C and alkylated with 15 mM iodoacetamide for 30 min at RT in the dark. The proteins on-bead were digested overnight at RT with 1 ug trypsin (Promega, V5113). The digested supernatant was acidified with 10% TFA to a pH of 2 and desalted with 10 mg StrataX solid phase extraction columns (Phenomenex), then dried under vacuum using a SpeedVac (Thermo Scientific) and stored at −80°C until MS analysis.

Samples were resuspended in 0.2% formic acid and subjected to LC-MS analysis. LC separation was performed using the Thermo Ultimate 3000 RSLCnano system. A 15 cm EASY-Spray™ PepMap™ RSLC C18 column (150 mm × 75 μm, 3 μm) was used at 300 nL/min flow rate with a 90 min gradient using mobile phase A consisting of 0.1% formic acid in H2O, and mobile phase B consisting of 0.1% formic acid in ACN/H2O (80/20, v/v). EASY-Spray source was used and temperature was at 35°C. Each sample run was held at 4.0% B for 5 min and increased to 50% B over 65 min, followed by 8 min at 95% B and back to 4% B for equilibration for 10 min. An Acclaim PepMap C18 HPLC trap column (20 mm × 75 μm, 3 μm) was used for sample loading. MS detection was performed with Thermo Exploris 240 Orbitrap mass spectrometer in positive mode. The source voltage was set to 1.8 kV, ion transfer tube temperature was set to 275°C, RF lens was at 70%. Full MS spectra were acquired from m/z 350 to 1400 at the Orbitrap resolution of 60000, with the normalized AGC target of 300% (3E6). Data-dependent acquisition (DDA) was performed for the top 20 precursor ions with the charge state of 2-6 and an isolated width of 2. Intensity threshold was 5E3. Dynamic exclusion was 30 s with the exclusion of isotopes. Other settings for DDA include Orbitrap resolution of 15000 and HCD collision energy of 30%.

Raw files were analyzed by SequestHT Search Engine incorporated in Proteome Discoverer v.2.5.0.400 software against human databases downloaded from Uniprot. Label-free quantification was enabled in the searches. The resulting data was analyzed by Perseus v 1.6.15.0 software ^45^.

#### Seahorse Analysis

HEK293 WT and Δ*COQ4* cells were split, and 4 x 10^5^ cells were plated per well in 6-well plates and allowed to grow overnight. On day two, cells were transiently transfected with a mix of 2 µg plasmid (pcDNA3.1 GFP-FLAG, pcDNA3.1 hCOQ4-FLAG WT, or pcDNA3.1 hCOQ4-FLAG D164A), 7.2 µg linear polyethylenimine (PEI, PolySciences), and 200 µL Opti-MEM (LifeTechnologies). On day 3, 40,000 cells per well were plated to Seahorse eXF24 plates coated with Cell-Tak (Corning) and allowed to adhere to the plate overnight in DMEM supplemented with 10% FBS and 1× penicillin– streptomycin. At 48 h post-transfection, and immediately before the Seahorse run, the medium was aspirated, cells were washed with dPBS, and medium was replaced with Seahorse XF DMEM Medium, pH 7.4 (Agilent no. 103575-100) supplemented with 10 mM glucose, 1 mM pyruvate and 2 mM glutamine. Oxygen-consumption rates (OCR) and extracellular acidification rate was monitored on a Seahorse eXF24 basally and in the presence of a Seahorse XF Cell Mito Stress Test (Agilent no. 103015-100). For the Stress Test, cells were treated with oligomycin (1 μM final concentration), FCCP (1 μM final concentration) and rotenone and antimycin A (0.5 μM final concentration). To normalize the cell number per well, cells were then fixed with 1% glutaraldehyde and stained with 1.5% crystal violet, and, after release of the stain with 10% acetic acid, each well was read at an absorbance of 590 nm. Data were exported from the Wave software (version 2.6.0) and analyzed using Prism (n = 5, error bars represent standard deviation).

### Quantification and statistical analysis

See each individual method and figures’ legends for the associated statistical analysis. The majority of *p* values in this report were calculated using an unpaired, one-sided, Student’s t test. In all cases, *n* represents independent replicates of an experiment and error bars represent standard deviation.

## SUPPLEMENTAL INFORMATION TITLES AND LEGENDS

Figure S1-S4, Table S1, Key Resources table

## REFERENCES

1. Baschiera, E., Sorrentino, U., Calderan, C., Desbats, M.A., and Salviati, L. (2021). The multiple roles of coenzyme Q in cellular homeostasis and their relevance for the pathogenesis of coenzyme Q deficiency. Free Radical Biology and Medicine 166, 277–286. 10.1016/j.freeradbiomed.2021.02.039.

2. Doimo, M., Desbats, M.A., Cerqua, C., Cassina, M., Trevisson, E., and Salviati, L. (2014). Genetics of coenzyme q10 deficiency. Mol Syndromol 5, 156–162. 10.1159/000362826.

3. Guerra, R.M., and Pagliarini, D.J. (2023). Coenzyme Q biochemistry and biosynthesis. Trends Biochem Sci 48, 463–476. 10.1016/j.tibs.2022.12.006.

4. Alcázar-Fabra, M., Rodríguez-Sánchez, F., Trevisson, E., and Brea-Calvo, G. (2021). Primary Coenzyme Q deficiencies: A literature review and online platform of clinical features to uncover genotype-phenotype correlations. Free Radic Biol Med 167, 141–180. 10.1016/j.freeradbiomed.2021.02.046.

5. Desbats, M.A., Morbidoni, V., Silic-Benussi, M., Doimo, M., Ciminale, V., Cassina, M., Sacconi, S., Hirano, M., Basso, G., Pierrel, F., et al. (2016). The COQ2 genotype predicts the severity of coenzyme Q10 deficiency. Human molecular genetics 25, 4256–4265. 10.1093/hmg/ddw257.

6. Stefely, J.A., and Pagliarini, D.J. (2017). Biochemistry of Mitochondrial Coenzyme Q Biosynthesis. Trends in biochemical sciences 42, 824–843. 10.1016/j.tibs.2017.06.008.

7. Ozeir, M., Muhlenhoff, U., Webert, H., Lill, R., Fontecave, M., and Pierrel, F. (2011). Coenzyme Q biosynthesis: Coq6 Is required for the C5-hydroxylation reaction and substrate analogs rescue Coq6 deficiency. Chem. Biol. 18, 1134–1142. 10.1016/j.chembiol.2011.07.008.

8. Acosta Lopez, M.J., Trevisson, E., Canton, M., Vazquez-Fonseca, L., Morbidoni, V., Baschiera, E., Frasson, C., Pelosi, L., Rascalou, B., Desbats, M.A., et al. (2019). Vanillic Acid Restores Coenzyme Q Biosynthesis and ATP Production in Human Cells Lacking COQ6. Oxidative medicine and cellular longevity 2019, 3904905. 10.1155/2019/3904905.

9. Wang, Y., and Hekimi, S. (2019). The Complexity of Making Ubiquinone. Trends in endocrinology and metabolism: TEM. 10.1016/j.tem.2019.08.009.

10. Marshall, S.A., Payne, K.A.P., and Leys, D. (2017). The UbiX-UbiD system: The biosynthesis and use of prenylated flavin (prFMN). Arch. Biochem. Biophys. 632, 209–221. 10.1016/j.abb.2017.07.014.

11. Hajj Chehade, M., Pelosi, L., Fyfe, C.D., Loiseau, L., Rascalou, B., Brugiere, S., Kazemzadeh, K., Vo, C.D., Ciccone, L., Aussel, L., et al. (2019). A Soluble Metabolon Synthesizes the Isoprenoid Lipid Ubiquinone. Cell chemical biology 26, 482–492 e7. 10.1016/j.chembiol.2018.12.001.

12. Floyd, B.J., Wilkerson, E.M., Veling, M.T., Minogue, C.E., Xia, C., Beebe, E.T., Wrobel, R.L., Cho, H., Kremer, L.S., Alston, C.L., et al. (2016). Mitochondrial Protein Interaction Mapping Identifies Regulators of Respiratory Chain Function. Molecular cell 63, 621–632. 10.1016/j.molcel.2016.06.033.

13. Hughes, B.G., Harrison, P.M., and Hekimi, S. (2017). Estimating the occurrence of primary ubiquinone deficiency by analysis of large-scale sequencing data. Scientific reports 7, 17744. 10.1038/s41598-017-17564-y.

14. Brea-Calvo, G., Haack, T.B., Karall, D., Ohtake, A., Invernizzi, F., Carrozzo, R., Kremer, L., Dusi, S., Fauth, C., Scholl-Burgi, S., et al. (2015). COQ4 mutations cause a broad spectrum of mitochondrial disorders associated with CoQ10 deficiency. American journal of human genetics 96, 309–317. 10.1016/j.ajhg.2014.12.023.

15. Mero, S., Salviati, L., Leuzzi, V., Rubegni, A., Calderan, C., Nardecchia, F., Galatolo, D., Desbats, M.A., Naef, V., Gemignani, F., et al. (2021). New pathogenic variants in COQ4 cause ataxia and neurodevelopmental disorder without detectable CoQ10 deficiency in muscle or skin fibroblasts. J Neurol 268, 3381–3389. 10.1007/s00415-021-10509-6.

16. Jurkute, N., Cancellieri, F., Pohl, L., Li, C.H.Z., Heaton, R.A., Reurink, J., Bellingham, J., Quinodoz, M., Yioti, G., Stefaniotou, M., et al. (2022). Biallelic variants in coenzyme Q10 biosynthesis pathway genes cause a retinitis pigmentosa phenotype. NPJ Genom Med 7, 60. 10.1038/s41525-022-00330-z.

17. 17. Salviati, L., Trevisson, E., Agosto, C., Doimo, M., and Navas, P. (1993). Primary Coenzyme Q10 Deficiency Overview. In GeneReviews®, M. P. Adam, J. Feldman, G. M. Mirzaa, R. A. Pagon, S. E. Wallace, L. J. Bean, K. W. Gripp, and A. Amemiya, eds. (University of Washington, Seattle).

18. Marbois, B., Gin, P., Gulmezian, M., and Clarke, C.F. (2009). The yeast Coq4 polypeptide organizes a mitochondrial protein complex essential for coenzyme Q biosynthesis. Biochim Biophys Acta 1791, 69–75. 10.1016/j.bbalip.2008.10.006.

19. Tran, U.C., and Clarke, C.F. (2007). Endogenous synthesis of coenzyme Q in eukaryotes. Mitochondrion 7, S62–71. 10.1016/j.mito.2007.03.007.

20. Kawamukai, M. (2016). Biosynthesis of coenzyme Q in eukaryotes. Biosci. Biotechnol. Biochem. 80, 23–33. 10.1080/09168451.2015.1065172.

21. Giorgio, V., Schiavone, M., Galber, C., Carini, M., Da Ros, T., Petronilli, V., Argenton, F., Carelli, V., Acosta Lopez, M.J., Salviati, L., et al. (2018). The idebenone metabolite QS10 restores electron transfer in complex I and coenzyme Q defects. Biochim Biophys Acta Bioenerg 1859, 901–908. 10.1016/j.bbabio.2018.04.006.

22. Casarin, A., Jimenez-Ortega, J.C., Trevisson, E., Pertegato, V., Doimo, M., Ferrero-Gomez, M.L., Abbadi, S., Artuch, R., Quinzii, C., Hirano, M., et al. (2008). Functional characterization of human COQ4, a gene required for Coenzyme Q(10) biosynthesis. Biochem. Biophys. Res. Commun. 372, 35–39. 10.1016/j.bbrc.2008.04.172.

23. Marbois, B., Gin, P., Faull, K.F., Poon, W.W., Lee, P.T., Strahan, J., Shepherd, J.N., and Clarke, C.F. (2005). Coq3 and Coq4 define a polypeptide complex in yeast mitochondria for the biosynthesis of coenzyme Q. J Biol Chem 280, 20231–20238. 10.1074/jbc.M501315200.

24. Zhang, H.T., and Javor, G.T. (2000). Identification of the ubiD gene on the Escherichia coli chromosome. J. Bacteriol. 182, 6243–6246. 10.1128/jb.182.21.6243-6246.2000.

25. Pelosi, L., Ducluzeau, A.L., Loiseau, L., Barras, F., Schneider, D., Junier, I., and Pierrel, F. (2016). Evolution of Ubiquinone Biosynthesis: Multiple Proteobacterial Enzymes with Various Regioselectivities To Catalyze Three Contiguous Aromatic Hydroxylation Reactions. mSystems 1, e00091–16. 10.1128/mSystems.00091-16.

26. Collins, M.D., and Jones, D. (1981). Distribution of isoprenoid quinone structural types in bacteria and their taxonomic implication. Microbiol Rev 45, 316–354.

27. Burgardt, A., Moustafa, A., Persicke, M., Spross, J., Patschkowski, T., Risse, J.M., Peters-Wendisch, P., Lee, J.-H., and Wendisch, V.F. (2021). Coenzyme Q(10) Biosynthesis Established in the Non-Ubiquinone Containing Corynebacterium glutamicum by Metabolic Engineering. Front. Bioeng. Biotechnol. 9, 650961. 10.3389/fbioe.2021.650961.

28. Burgardt, A., Pelosi, L., Chehade, M.H., Wendisch, V.F., and Pierrel, F. (2022). Rational Engineering of Non-Ubiquinone Containing Corynebacterium glutamicum for Enhanced Coenzyme Q10 Production. Metabolites 12, 428. 10.3390/metabo12050428.

29. 29. Yang, Y.L., Zhou, H., Du, G., Feng, K.N., Feng, T., Fu, X.L., Liu, J.K., and Zeng, Y. (2016). A Monooxygenase from Boreostereum vibrans Catalyzes Oxidative Decarboxylation in a Divergent Vibralactone Biosynthesis Pathway. Angewandte Chemie. 10.1002/anie.201510928.

30. Herebian, D., Seibt, A., Smits, S.H.J., Bunning, G., Freyer, C., Prokisch, H., Karall, D., Wredenberg, A., Wedell, A., Lopez, L.C., et al. (2017). Detection of 6-demethoxyubiquinone in CoQ10 deficiency disorders: Insights into enzyme interactions and identification of potential therapeutics. Molecular genetics and metabolism. 10.1016/j.ymgme.2017.05.012.

31. Pierrel, F. (2017). Impact of Chemical Analogs of 4-Hydroxybenzoic Acid on Coenzyme Q Biosynthesis: From Inhibition to Bypass of Coenzyme Q Deficiency. Frontiers in Physiology 8. 10.3389/fphys.2017.00436.

32. Nagatani, H., Mae, Y., Konishi, M., Matsuzaki, M., Kita, K., Daldal, F., and Sakamoto, K. (2023). UbiN, a novel Rhodobacter capsulatus decarboxylative hydroxylase involved in aerobic ubiquinone biosynthesis. FEBS Open Bio. 10.1002/2211-5463.13707.

33. Ozeir, M., Pelosi, L., Ismail, A., Mellot-Draznieks, C., Fontecave, M., and Pierrel, F. (2015). Coq6 Is Responsible for the C4-deamination Reaction in Coenzyme Q Biosynthesis in Saccharomyces cerevisiae. The Journal of biological chemistry 290, 24140–24151. 10.1074/jbc.M115.675744.

34. Schenk, G., Mitić, N., Gahan, L.R., Ollis, D.L., McGeary, R.P., and Guddat, L.W. (2012). Binuclear Metallohydrolases: Complex Mechanistic Strategies for a Simple Chemical Reaction. Acc. Chem. Res. 45, 1593–1603. 10.1021/ar300067g.

35. Cerqua, C., Casarin, A., Pierrel, F., Vazquez Fonseca, L., Viola, G., Salviati, L., and Trevisson, E. (2019). Vitamin K2 cannot substitute Coenzyme Q10 as electron carrier in the mitochondrial respiratory chain of mammalian cells. Scientific reports 9, 6553. 10.1038/s41598-019-43014-y.

36. Burke, D., Dawson, D., and Stearns, T. (2000). In Methods in Yeast Genetics (Cold Spring Harbor Laboratory Press, Plainview, NY).

37. Bott, L.E., Michael ed. (2005). Handbook of Corynebacterium glutamicum (CRC Press) 10.1201/9781420039696.

38. Fukasawa, Y., Tsuji, J., Fu, S.-C., Tomii, K., Horton, P., and Imai, K. (2015). MitoFates: improved prediction of mitochondrial targeting sequences and their cleavage sites. Mol Cell Proteomics 14, 1113–1126. 10.1074/mcp.M114.043083.

39. Haupka, C. (2020). Codon harmonization.

40. Kirchner, O., and Tauch, A. (2003). Tools for genetic engineering in the amino acid-producing bacterium Corynebacterium glutamicum. J Biotechnol 104, 287–299. 10.1016/s0168-1656(03)00148-2.

41. Cherepanov, P.P., and Wackernagel, W. (1995). Gene disruption in Escherichia coli: TcR and KmR cassettes with the option of Flp-catalyzed excision of the antibiotic-resistance determinant. Gene 158, 9–14. 10.1016/0378-1119(95)00193-a.

42. Thomason, L.C., Costantino, N., and Court, D.L. (2007). E. coli genome manipulation by P1 transduction. Curr Protoc Mol Biol Chapter 1, Unit 1.17. 10.1002/0471142727.mb0117s79.

43. Pelosi, L., Vo, C.D., Abby, S.S., Loiseau, L., Rascalou, B., Hajj Chehade, M., Faivre, B., Gousse, M., Chenal, C., Touati, N., et al. (2019). Ubiquinone Biosynthesis over the Entire O2 Range: Characterization of a Conserved O2-Independent Pathway. mBio 10, e01319–19. 10.1128/mBio.01319-19.

44. Kazemzadeh, K., Pelosi, L., Chenal, C., Chobert, S.-C., Hajj Chehade, M., Jullien, M., Flandrin, L., Schmitt, W., He, Q., Bouvet, E., et al. (2023). Diversification of ubiquinone biosynthesis via gene duplications, transfers, losses, and parallel evolution. Mol Biol Evol, msad219. 10.1093/molbev/msad219.

45. Tyanova, S., and Cox, J. (2018). Perseus: A Bioinformatics Platform for Integrative Analysis of Proteomics Data in Cancer Research. Methods Mol Biol 1711, 133–148. 10.1007/978-1-4939-7493-1_7.

