## Supplemental Figures 1-4 and tables for "COQ4 is required for the oxidative decarboxylation of the C1 carbon of Coenzyme Q in eukaryotic cells"

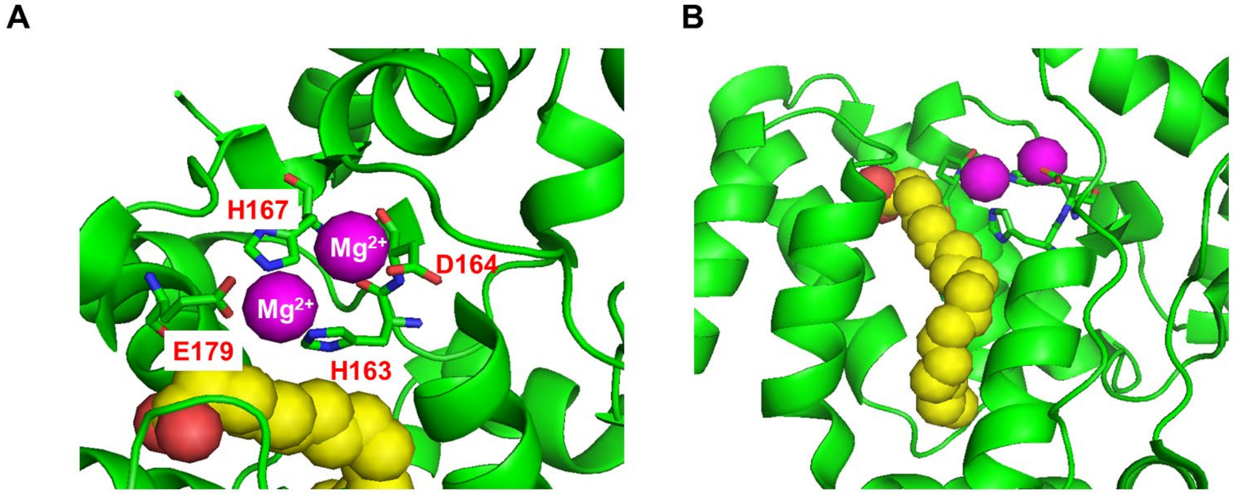

**Figure S1**: A) Structure of the metal binding domain of the COQ4 homolog Alr8543 from Nostoc sp. PCC 7120 (PDB: 6E12) visualized with PyMol. Highlighted are the two $Mg^{2+}$ ions, and the residues corresponding to histidine 163, aspartate 164, histidine 167, and glutamate 179 of the human protein which define the HDxxH(x)11E motif. B) The hydrophobic tunnel containing an oleic acid molecule.

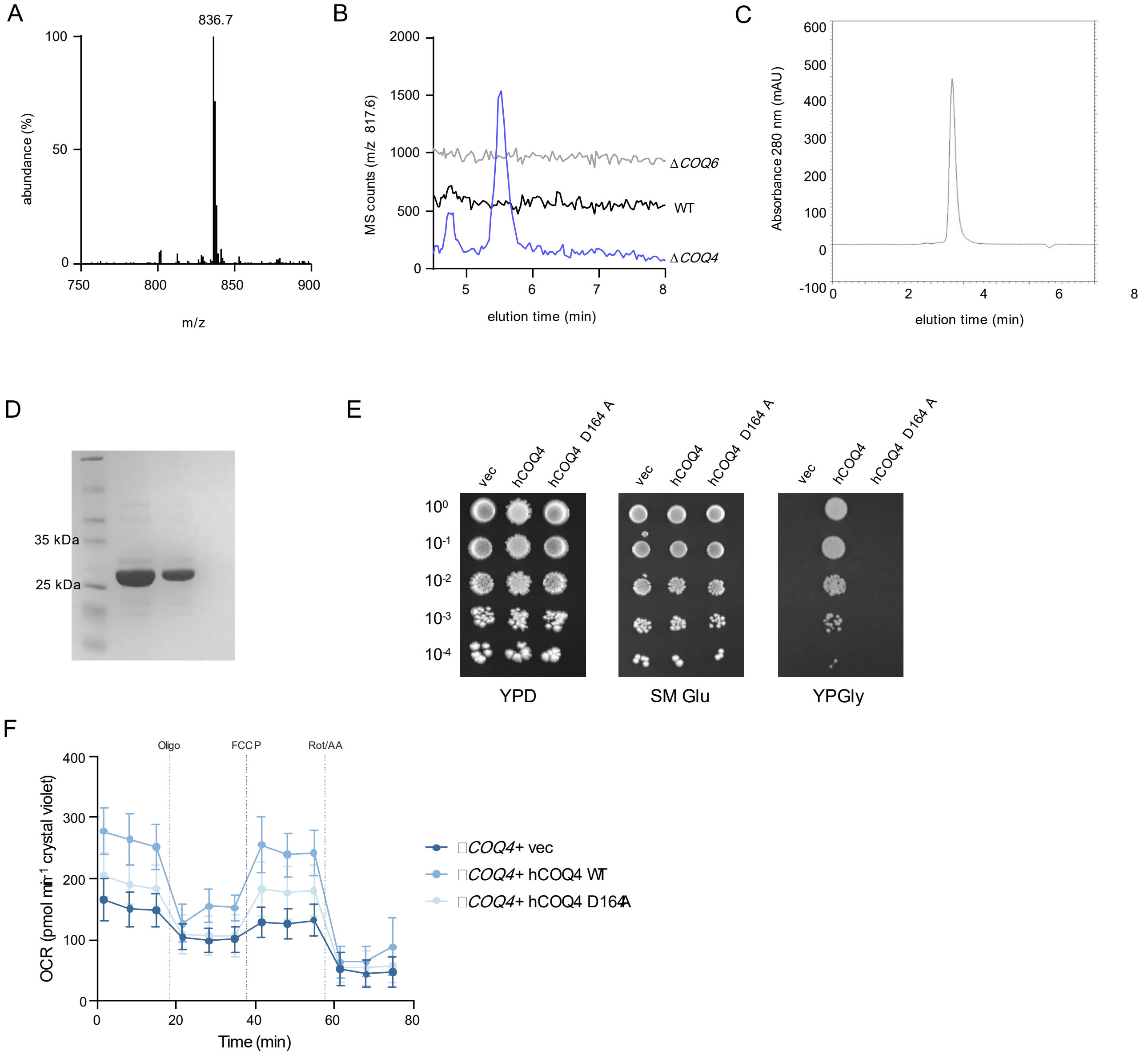

**Figure S2**: A) Mass spectrum of the compound eluting at 5.6 min in Δ*COQ4* extracts (see Fig 1B). B) Single ion monitoring in negative mode (m/z 817.6, M$^-$ for $_{10}$P-HB) in HPLC-ECD-mass spectrometry analyses. The lipid extracts from human WT, Δ*COQ4* and Δ*COQ6* HEK293 cells (0.2 mg protein) were analyzed with the precolumn electrode in oxidizing mode. Results representative of 3 biological triplicates. C) Elution profile of recombinant hCOQ4 from the Superdez Yarra SEC-2000 gel filtration column (see methods) monitored at 280 nm. D) Coomassie stained SDS-PAGE of the collected fraction and of the same sample diluted 1:2. The molecular ladder used is SHARPMASS VI (Euroclone). E) Plate dilution assays of yeast *W303*Δ*COQ4* overexpressing yCOQ8 from a high copy vector and transformed with either wild type human COQ4, the D164A allele, or the empty vector (vec). The cells were grown for 5 days at 30°C. The overexpression of yCOQ8 is necessary to stabilize complex Q in the *W303*Δ*COQ4* cells, as in any yeast strain knocked-out for *coq* genes [1]. F) Oxygen consumption rates from a Seahorse mitochondria stress test of HEK293 Δ*COQ4* cells expressing GFP control empty vector, hCOQ4 WT, or hCOQ4 D164A, shown as mean ± SD (n=5); Oligo, oligomycin; FCCP, carbonyl cyanide 4-(trifluoromethoxy)phenylhydrazone; Rot + AA, rotenone + antimycin A.

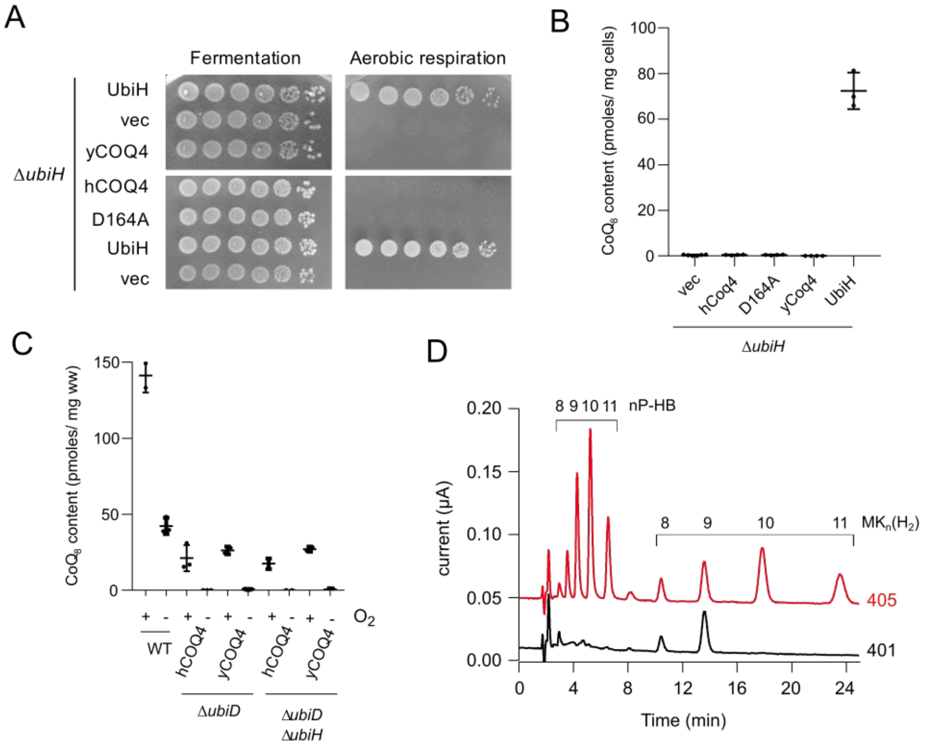

**Figure S3:** A) Plate-growth assay of serial dilutions of *E. coli* Δ*ubiH* cells containing empty vectors (vec) or vectors expressing UbiH, yeast Coq4 (yCOQ4), human COQ4 (hCOQ4) or the human COQ4 D164A allele (D164A). B) $CoQ_8$ content of Δ*ubiH* cells containing vectors described in A, after growth in LB medium. C) $CoQ_8$ content of Δ*ubiD* and Δ*ubiD* Δ*ubiH* cells expressing hCOQ4 or yCOQ4, after growth in LB medium under aerobic ($+O_2$) or anaerobic ($-O_2$) conditions. D) Electrochromatograms obtained by HPLC-ECD-MS analysis of lipid extracts from *C. glutamicum* strains 401 and 405. See Brugardt et al [2] for the identification of the compounds nP-HB and $MK_n(H_2)$ by mass spectrometry.

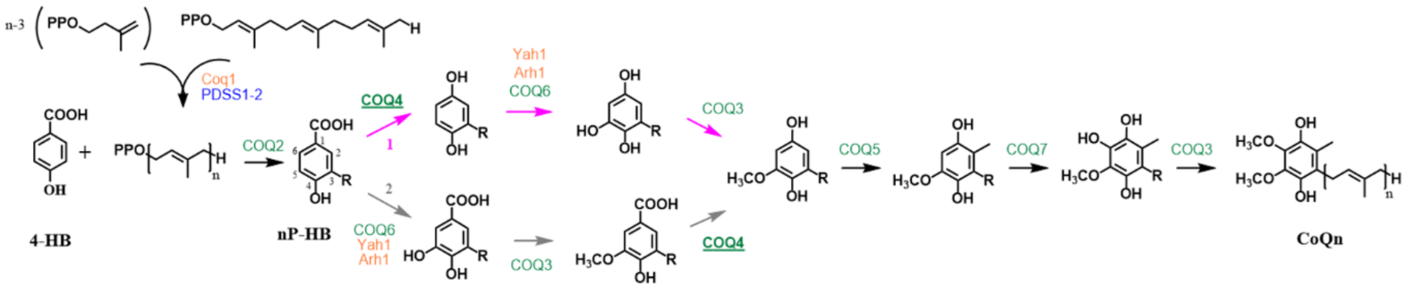

**Figure S4:** Proposed new biosynthetic pathway of CoQ in eukaryotes. See Figure 1 for additional details and comparison. Proteins shared by *S. cerevisiae* and humans are in green, and those specific to *S. cerevisiae* or humans are in orange and blue, respectively. Based on our results, we propose that position C1 of polyprenyl-4-HB (nP-HB) undergoes an oxidative decarboxylation catalyzed by COQ4 and that position C5 is later modified by COQ6 and COQ3 (path 1 in pink). Alternatively, COQ6 and COQ3 may act before COQ4 (path 2 in grey). The polyprenyl chain (n=6 for *S. cerevisiae*, n=10 for humans) is depicted by R on all intermediates derived from nP-HB.

**Table S1. Oligonucleotides used in the study**

| Name | Sequence (5'->3') | Application |
|---|---|---|
| hCOQ4_BAC_NheI_F | CTTCTGCTAGCAGGAGGAAATCACCATGAAATGCCCCTCCGGGCTAGG | Cloning of pBAD-hCOQ4 |
| hCOQ4_HindIII_R | CTTCTAAGCTTGCTCAGGAGCTCAGGCCAA | Cloning of pBAD-hCOQ4 and pET21a-hCOQ4 |
| hCOQ4_HisT_NdeI_For | CTTCTCATATGCATCACCATCACCATCACATTGATGACGACGACAAGGCGGCAGAAATGCCCCTCCGGGC | Cloning of pET21a-hCOQ4 |
| D164A_F | CGGGAGGTGCACGCCATGCTTCACACC | Mutagenesis |
| D164A_R | GGTGTGAAGCATGGCGTGCACCTCCCG | |
| UbiHprom5 | ACGCCCATCGTCCTTTCTTT | Verification of strain *ΔubiDΔubiH* |
| UbiHterm3 | CAGTTGTGGTGGTGCATTCG | |
| yCOQ4_Sac_Fw | CCTACCgagctcATGTTGAGGTTATCTTTACTGAG | Cloning of pTrc-yCOQ4 |
| yCOQ4_HIII_Rv | CTGGAGaagcttTCATGCTGGAGTCGTGGCTCG | |
| UbiH5-EcoRI | cctaagaattcATGAGCGTAATCATCGTCGG | Cloning of pTrc-UbiH |
| UbiH3-HindIII | ttaggaagcttTCAACGCGCCACCCAACCG | |
| ubiX-fw | TTCGAGCTCGGTACCCGGGGATCCTGAAAGGAGGCCCTTCAGATGAAACGACTCATTGTAGGCAT | Cloning of pEC-XT99A-*ubiDX* |
| ubiX-rv | TTGCATGCCTGCAGGTCGACTCTAGTTATGCGCCCTGCCAGCGG | |
| yCOQ4-CG-fw | ATGGAATTCGAGCTCGGTACCCGGGGAAAGGAGGCCCTTCAGATGCTGCGTCTGTCCCTGCTAC | Cloning of pEC-XT99A-yCOQ4 |
| yCOQ4-CG-rv | GCCTGCAGGTCGACTCTAGAGGATCTTATGCTGGGGTCGTCGCCCTC | |
| hCOQ4_D164A_F | GAGGTGCACGcCATGCTTCAC | Mutagenesis |
| hCOQ4_D164A_R | CCGGTACCGCTGAATCAC | |
| yCOQ3 HindIII F | AATAATAAGCTTATGAAGAGCACAGATGCTTCTGAA | Cloning of pET28b NΔ31 yCOQ3 |
| yCOQ3 XhoI R | ATTATTCTCGAGTCAATTCAGTCTCTGAATAGCCAT | |
| yCOQ5 HindIII F | AATAATAAGCTTATGAAAGAAGAAGAAGTTAATAGT | Cloning of pET28b NΔ30 yCOQ5 |
| yCOQ5 XhoI R | ATTATTCTCGAGTTAAACTTTAATGCCCCAATGGAT | |
| yCOQ9 HindIII F | AATAATAAGCTTATGTATCATTCGAACCCTATCGAA | Cloning of pET28b NΔ16 yCOQ9 |
| yCOQ9 XhoI R | ATTATTCTCGAGTTAACCCCTAACTAATTGAGATTT | |
| yCOQ6 Gibson F | CCAGGATCCGGTCTCAAGGTATGTTCTTCAGCAAGGTCATG | Cloning of pET28b yCOQ6 |
| yCOQ6 Gibson R | GCTCGAATTCTCTAGATTATCACTTCTCGTTTCCGCCCAG | |
| pET28b Gibson F | CTGGGCGGAAACGAGAAGTGATAATCTAGAGAATTCGAGC | |
| pET28b Gibson R | CATGACCTTGCTGAAGAACATACCTTGAGACCGGATCCTGG | |

### KEY RESOURCES TABLE

| REAGENT or RESOURCE | SOURCE | IDENTIFIER |
|---|---|---|
| Antibodies | | |
| Not relevant | | |
| Bacterial and virus strains | | |
| *E. coli* Mach1 competent cells | Thermo Scientific™ | Cat#C862003 |
| *E. coli* BL21 (DE3) competent cells | Sigma Aldrich (Merck) | Cat#69450-M |
| *E. coli* DH5α competent cells | Thermo Scientific™ | Cat#18265017 |
| *E. coli* MG1655 | Lab stock, Pelosi et al. [26] | N/A |
| *E. coli* Δ*ubiD::cat* | Pelosi et al. [26] | N/A |
| *E. coli* Δ*ubiD* | This paper | N/A |
| *E. coli* Δ*ubiD* Δ*ubiH* | This paper | N/A |
| *E. coli* Δ*ubiH* | Pelosi et al. [26] | N/A |
| *C. glutamicum* UBI400 | Burgardt et al. [28] | N/A |
| *C. glutamicum* 401 | This paper | N/A |
| *C. glutamicum* 405 | This paper | N/A |
| Biological samples | | |
| Not relevant | | |
| Chemicals, peptides, and recombinant proteins | | |
| isopropyl-1-thio-β-D-galactopyranoside | Euroclone | Cat#EMR066005 |
| N-Lauroylsarcosine | Sigma Aldrich (Merck) | Cat#L9150 |
| Anhydrotetracycline (hydrochloride) | biomol | Cat#Cay10009542-100 |
| $^{13}C_7$-4HB | Sigma Aldrich (Merck) | Cat#606472 |
| Digitonin | MilliporeSigma | Cat#D141-500MG |
| Coenzyme Q6 | Avanti Lipids | Cat#900150O-1mg |
| Coenzyme Q8 | Avanti Lipids | Cat#900151P-1mg |
| Trypsin | Promega | Cat#V5113 |
| Polyethylenimine | Polysciences | Cat#19850 |
| CellTak | Corning | Cat#354240 |
| cOmplete protease inhibitor cocktail | MilliporeSigma (Roche) | Cat#11697498001 |
| Critical commercial assays | | |
| QuikChange lightning site-directed mutagenesis kit | Agilent | Cat#210518 |
| Seahorse XFe24 FluxPak Mini | Agilent | Cat#102342-100 |
| Seahorse XF Cell Mito Stress Test | Agilent | Cat#103015-100 |
| Seahorse XF DMEM Medium | Agilent | Cat#103575-100 |
| NEBuilder® HiFi DNA Assembly Cloning Kit | New England Biolabs | Cat#E5520S |
| Deposited data | | |
| Affinity enrichment mass spectrometry data | MassiVE | MSV000093201 |
| Experimental models: Cell lines | | |
| HEK 293 ΔCOQ4 | Cerqua et al.[35] | N/A |

| | | |
|---|---|---|
| HEK 293 ΔCOQ6 | Acosta Lopez et al.[8] | N/A |
| Human primary fibroblasts carrying the COQ4 mutation c.577C>T/p.Pro193Ser in trans with c.718C>T/p.Arg240Cys | Mero et al.[15] | N/A |
| Experimental models: Organisms/strains | | |
| *S. cerevisiae* W303 Δ*coq4* | Mero et al.[15] | N/A |
| Oligonucleotides | | |
| Oligonucleotides used in this study are listed in Table S1 | | |
| Recombinant DNA | | |
| Plasmid: pRG_Duet2 | Burgardt et al. [27] | Genbank: MK130721 |
| Plasmid: pRG_Duet2-*ddsA*$_{Pd}$-*ubiA*$_{Ec}$ | Burgardt et al. [27] | N/A |
| Plasmid: pEC-XT99A | Kirchner and Tauch [40] | Genbank: AY219684 |
| Plasmid: pEC-XT99A-*ubiD* | Burgardt et al. [27] | N/A |
| Plasmid: pEC-XT99A-*ubiDX* | This paper | N/A |
| Plasmid: pEC-XT99A-yCOQ4 | This paper | N/A |
| Gene synthesis: COQ4 from *S. cerevisiae* codon-harmonized to *C. glutamicum* ATCC 13032 | This paper | N/A |
| Plasmid: pET21a | Novagen | Cat#69740-3 |
| Plasmid: pBAD24 | ATTC | Cat#87399 |
| Plasmid: pBAD-hCOQ4 | This paper | N/A |
| Plasmid: pET21a-hCOQ4 | This paper | N/A |
| Plasmid: pTrc-yCOQ4 | This paper | N/A |
| Plasmid: pTrc-UbiH | This paper | N/A |
| Plasmid: pcDNA3.1 GFP-FLAG | Floyd et al. [12] | N/A |
| Plasmid: pcDNA3.1 hCOQ4-FLAG | Floyd et al. [12] | N/A |
| Plasmid: pcDNA3.1 hCOQ4-FLAG D164A | This paper | N/A |
| Plasmid: pET28b NΔ31 yCOQ3 | This paper | N/A |
| Plasmid: pET28b NΔ30 yCOQ5 | This paper | N/A |
| Plasmid: pET28b NΔ16 yCOQ9 | This paper | N/A |
| Plasmid: pET28b yCOQ6 | This paper | N/A |
| Gene synthesis: COQ6 from *S. cerevisiae* codon-harmonized to *E. coli*. | This paper | N/A |
| Plasmid: pRS423-yCOQ8 | Ozeir et al. [33] | N/A |
| Plasmid: pCM189_COQ4 | Mero et al.[15] | N/A |
| Plasmid: pCM189_COQ4$^{D164A}$ | This paper | N/A |
| Software and algorithms | | |
| TraceFinder 5.1 | ThermoFisher | N/A |
| XCalibur 4.4 | ThermoFisher | N/A |
| Proteome Discoverer v2.5.0.400 | ThermoFisher | N/A |
| Perseus v1.6.15.0 | Tyanova and Cox [45] | N/A |
| Prism v9.4.1 | GraphPad | N/A |
| MitoFates | Fukasawa et al. [38] | N/A |
| Wave v2.6.0 | Agilent | N/A |
| Other | | |
| SnakeSkin Dialysis Tubing, 3,500 MWCO | Thermo Scientific™ | Cat#68100 |
| Superdez Yarra SEC-2000 column | Phenomenex | Cat#00H-4512 |

| | | |
|---|---|---|
| Ni affinity column HisTrap HP, 5ml | Cytiva | Cat#17524801 |
| BD GasPak™ EZ Gas Generating Systems | Fisher Scientific | Cat#260001 |
| Anti-FLAG® M2 Magnetic Beads | Sigma | Cat#M8823 |
| StrataX solid phase extraction columns | Phenomenex | Cat# 8B-S100-AAK |

### Supplemental references

1. Xie, L.X., Ozeir, M., Tang, J.Y., Chen, J.Y., Kieffer-Jaquinod, S., Fontecave, M., Clarke, C.F., and Pierrel, F. (2012). Over-expression of the Coq8 kinase in Saccharomyces cerevisiae coq null mutants allows for accumulation of diagnostic intermediates of the Coenzyme Q6 biosynthetic pathway. J Biol Chem *287*, 23571–23581. 10.1074/jbc.M112.360354.

2. Burgardt, A., Pelosi, L., Chehade, M.H., Wendisch, V.F., and Pierrel, F. (2022). Rational Engineering of Non-Ubiquinone Containing Corynebacterium glutamicum for Enhanced Coenzyme Q10 Production. Metabolites *12*, 428. 10.3390/metabo12050428.
